# Gsta4 controls apoptosis of differentiating adult oligodendrocytes during homeostasis and remyelination via the mitochondria-associated Fas/Casp8/Bid-axis

**DOI:** 10.1101/2020.05.26.111856

**Authors:** Karl E Carlström, Keying Zhu, Ewoud Ewing, Inge E Krabbendam, Robert A Harris, Ana Mendanha Falcão, Maja Jagodic, Gonçalo Castelo-Branco, Fredrik Piehl

## Abstract

Arrest of oligodendrocyte (OL) differentiation and remyelination following myelin damage in Multiple Sclerosis (MS) are associated with disease progression but the underlying mechanism are elusive. We show that Glutathione S-transferase 4α (Gsta4) is highly expressed during adult OL differentiation and that its loss prevents differentiation into myelinating OLs. Also, Gsta4 appeared to be a novel target for Clemastine, in clinical trial for MS. Over-expression of Gsta4 reduced the expression of Fas and activity along the mitochondria-associated Casp8/Bid-axis in adult pre-OLs from the corpus callosum, together promoting enhanced pre-OL survival during differentiation. The Gsta4-mediated input on apoptosis during adult OL differentiation was further verified in the LPC and EAE model, where Casp8 were reduced in pre-OLs with high Gsta4 expression in an immune response-independent fashion. Our results place Gsta4 as a key regulator of intrinsic mechanisms beneficial for OL differentiation and remyelination, and as a possible target for future MS therapies.

## Introduction

Oligodendrocyte precursor cells (OPC) and oligodendrocytes (OL) are abundant throughout the central nervous system (CNS). They serve important roles in preserving the functionality and integrity of neuronal network connections, including providing metabolic support and enabling axonal signal transmission along myelinated fibers ^1–4^. OLs are affected directly or indirectly in many CNS diseases, of which Multiple Sclerosis (MS) is one of the most widely studied ^5^. Despite findings of proliferating OPCs around lesions ^6,7^, OLs fail to remyelinate axons ^8–12^ which leads to neurodegeneration ^9,13–15^.

Previous studies have indicated that as many as 50% of pre-OLs undergo programmed cell death during development and 20-30% during post-natal conditions ^16,17^. Recent studies have also shown mechanisms of OLs loss during development ^18,19^, similar approached in post-natal animals and adult OLs are lacking.

Remyelination do occur in adulthood ^6, 20–23^, but the time interval within which the OLs should receive crucial support in order to survive is restricted ^24–26^.

The OPCs capacity to remyelinate is strongly diminished over-time ^27–29^ and differentiating cells are exposed to considerable cellular stress ^30–33^. Interestingly, mitochondrial function and the capacity to scavenge endogenous toxins are also diminished with ageing and disease-processes such as those present in MS ^34–36^. Notably, the activity of the primary sensor of cellular stress, Nuclear factor (erythroid-derived 2)-like 2 (Nrf2), is also diminished with ageing ^37,38^. Dysfunctional mitochondria and toxin scavenging induce lipid peroxidation (LPO) of unsaturated lipids in membranes, against which OLs are particularly vulnerable to ^39–42^. Upon LPO, reactive aldehydes, such as 4-Hydroxynonenal (4-HNE), will be generated and diffuse in the cytosol, unless scavenged by conjugation to glutathione. If not scavenged, 4-HNE will bind to macro-molecules and proteins, affecting their functions ^43,44^ and set a brake on several cellular processes ^45–47^.

In this study, we show that Glutathione S-transferase 4α (Gsta4) is highly expressed during OL differentiation. Interestingly, we also identify Gsta4 as novel target for the remyelinating drug Clemastine through Nrf2. Moreover, Gsta4 restricted apoptosis in pre-OLs via modulation of the Fas/Casp8 pathway thus facilitating a higher amount of OPC to generate myelinating OLs. Importantly, Gsta4 also promoted remyelination and ameliorated MS-like disease.

## Results

### Clemastine and dimethyl fumarate upregulate Gsta4 and promotes OL differentiation in a Gsta4-dependent manner

The transcription factor Nrf2 has been reported to be the main therapeutic target of DMF in several cell types, including OLs ^48–51^. Further, the main scavenger of 4-HNE, Gsta4, has been predicted to be a target of Nrf2 ^52,53^. We thus wanted to investigate whether a DMF-Nrf2-Gsta4 axis is operation in the oligodendrocyte lineage. As previously described by others ^49^, we observed that DMF contributed to OL maturation, as assessed by expression of myelin proteolipid protein (*Plp*) mRNA upon differentiation of rat OPCs *in vitro* (Fig.1a, Fig.S1a) Interestingly, we also observed that a drug formulation of clemastine, containing a fumarate moiety (clemastine-fumarate, CF) (Fig.1b), which has shown promising clinical effects in a trial of MS patients with chronic optic neuropathy ^54–56^, also increase *Plp* expression (Fig.1a). As expected, both DMF and CF caused nuclear activation of Nrf2 in OLs (Fig.1c). Notably, DMF and CF also induced the expression of *Gsta4* (Fig.1d). Importantly, *Gsta4* transcription was strongly increased in OLs during differentiation compared to proliferating OPCs (Fig.1e). DMF and CF mediated *Plp* expression was reduced in the presence of Gsta4-specific siRNA (siRNA-Gsta4) knockdown (Fig.1f, Fig.S1b). Thus, *Gsta4* acts downstream of DMF and CF to promote OL differentiation.

**Fig. 1.**
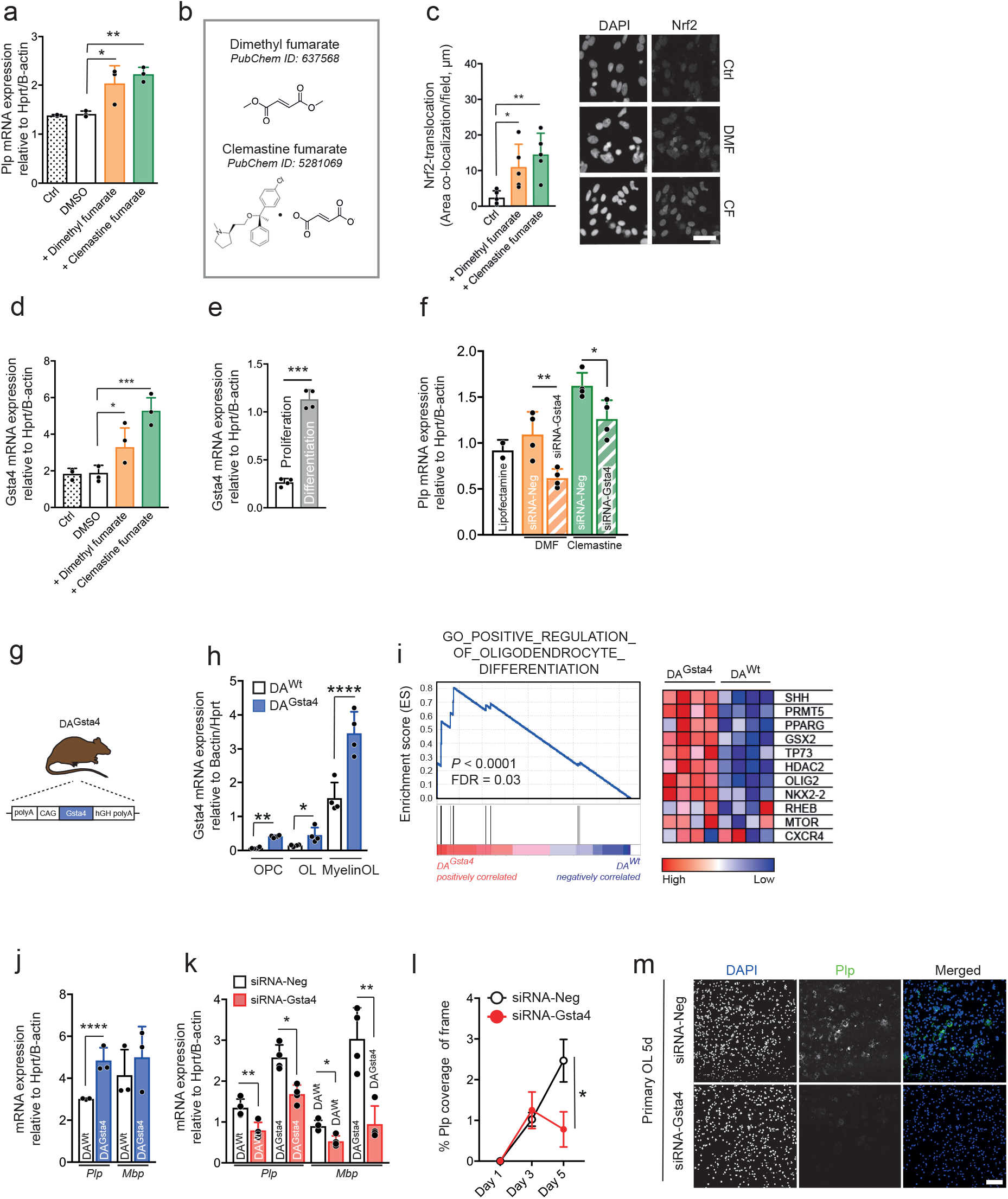
Clemastine and dimethyl fumarate upregulate Gsta4 and promotes OL differentiation in a Gsta4-dependet manner. (a) Transcription of *Plp* in primary OL cultures following stimulation for 48h with DMF (30μM) and CF (15μM) dissolved in DMSO (n=3). (b) Chemical structure of DMF and CF. (b) Nuclear activation of Nrf2 following stimulation for 24h with DMF (30μM) and CF (15μM) dissolved in DMSO (scale bar 50μm) (n=5). (d) Transcription of *Gsta4* in primary OL cultures following stimulation for 48h with DMF (30μM) and CF (15μM) dissolved in DMSO (n=3). (e) Transcription of *Gsta4* in proliferative OPCs compared to differentiated OLs *in vitro* (n=4). (f) Transcription of *Plp* in primary OL cultures following stimulation for 24h with DMF (30μM) and CF (15μM) dissolved in DMSO and lipofectamine-mediated knockdown of Gsta4 with siRNA-Gsta4 compared to scrambled sequence, siRNA-Neg (n=4). (g) Illustration of Dark Agouti (DA) rat, overexpressing Gsta4 (DA^Gsta4^). (h) *In vivo* transcription of *Gsta4* across the OL lineage in adult DA^Gsta4^ and DA^Wt^ (n=4). (i) Gene Set Enrichment Analysis (GSEA), analyzed with weighted enrichment statistics and ratio of classes for the metric as input parameters, on CNS bulk tissue (cortex, corpus callosum and hippocampus) from adult DA^Gsta4^ and DA^Wt^ (n=4). (j) Transcription of *Mbp* and *Plp* during differentiation of DA^Gsta4^ or DA^Wt^-derived OPCs (n=3). (k) Transcription of *Mbp* and *Plp* during differentiation of DA^Gsta4^ or DA^Wt^-derived OPCs in combination with Gsta4 knockdown with siRNA-Gsta4 or siRNA-Neg as a scrambled sequence (n=4). (l) Immunocytochemistry of Plp protein expression following Gsta4 knockdown with siRNA-Gsta4 (n=10) (m). Representative images at day 5, scale bar 50μm. (n) *In vitro* proliferation of DA^Wt^ or DA^Gsta4^-derived OPCs without (n=4) or with (n=6) application of Gsta4 knock-down with siRNA-Gsta4. (o) Representative contrast images of all four conditions, scale-bar 100μm. Graph show representative results from three (a, c, d, f, k, n) or two (e, h, j) independent experiments. All data are shown as mean and S.D. All statistical analyses were performed using one-way ANOVA. \**P*<0.05, \*\**P* <0.01, \*\*\**P*<0.001,\*\*\*\**P*<0.0001.

### Overexpression of Gsta4 in rats leads to increase expression of genes involved in OL differentiation

To study *in vivo* effects of Gsta4 we created a hemizygous rat (DA^Gsta4^), over-expressing *Gsta4* under a CAG promotor on a Dark Agouti (DA) background (Fig.1g). The DA^Gsta4^ strain displayed approximately a two-fold higher levels of *Gsta4* mRNA throughout the OL lineage compared to DA^Wt^ and the elevation was more pronounced in more mature cells (Fig.1h). DA^Gsta4^ also exhibited elevated *Gsta4* mRNA levels in additional tissues, including brain, spinal cord and spleen (data not shown). Nevertheless, DA^Gsta4^ showed no obvious phenotypic characteristics in terms of behavior, general health, spontaneous tumors, weight fertility or litter size as compared to wild type animals (DA^Wt^) (data not shown). The initial characterization of DA^Gsta4^ was performed with mRNA microarray of bulk tissue from adult (10-12 weeks) brain cortex, corpus callosum and hippocampus. Gene Set Enrichment Analysis (GSEA) on the mRNA indicated that transcripts involved in positive regulation inducing OL differentiation were significantly enriched in DA^Gsta4^ compared to DA^Wt^ (Fig.1i).

To assess if this translated into differences in OL differentiation and maturation markers, we determined the expression of *Plp* and myelin basic protein (*Mbp*) in primary oligodendrocyte cultures established from neonatal pups (Fig.S1a). Transcription of *Plp* were more prominent in DA^Gsta4^ OLs compared to DA^Wt^ when OPCs were cultured in differentiation media for 5 days (Fig.1j). Knock-down of *Gsta4* expression with siRNAs induced a reduction of *Plp* and *Mbp* mRNA upon differentiation of DA^Wt^ or DA^Gsta4^ OPCs (Fig.1k). Moreover, the production of Plp protein during differentiation was also decreased following addition of siRNA-Gsta4 compared to siRNA-Neg (Fig.1l,m, Fig.S1b). In addition, Gsta4 over-expression promoted enhanced proliferation of OPCs derived from neonatal pups compared to wild type conditions, whereas proliferation was reduced upon knock down with siRNA (Fig.1, n, o, Fig.S1a). Collectively, these findings highlight a beneficial impact of Gsta4 during OL lineage progression and in response to exposure remyelinating drugs.

### Gsta4 accelerates the transition of OPC to myelinating OLs by mitigating Casp8/Bid activity

The OL lineage was further evaluated in adult DA^Wt^ and DA^Gsta4^ rats as illustrated in Fig.2a. In corpus callosum (CC), adult DA^Gsta4^ was found to have a smaller pool of PDGFRα^+^ OPC (Fig.2b). Furthermore, flow cytometric analysis of CC and grey matter regions indicated in Fig.2a (*dotted line*) validated the histology finding where PDGFRα^+^ OPCs and O4^+^ OLs were significantly decreased in DA^Gsta4^ compared to DA^Wt^ whereas no strain-differences were evident in mature O1^+^ OLs (Fig.S1d,e). However, the pool of OPCs was found to be more proliferative since a larger proportion of the DA^Gsta4^ OPCs were positive for Ki67^+^ compared to DA^Wt^ (Fig.2c, d). We also analyzed whether Gsta4 was affecting adult OPC differentiation, by assessing EdU in O4^+^ and O1^+^ OLs, after administration of EdU in drinking water for 10 days and 6 weeks or after a single injection of EdU. The proportion of both double-positive O4^+^EdU^+^ and O1^+^EdU^+^ of all O4^+^ or O1^+^ were higher in DA^Gsta4^ at 10 days and 6 weeks (Fig.2e,f) supporting a faster differentiation in DA^Gsta4^. Overall, the total number of CC1^+^ OLs in the adult CC was not different between DA^Gsta4^ and DA^Wt^ (Fig.2g), suggesting a relative balance between of the effects of Gst4a on OL differentiation and OPC proliferation *in vivo*.

**Fig. 2.**
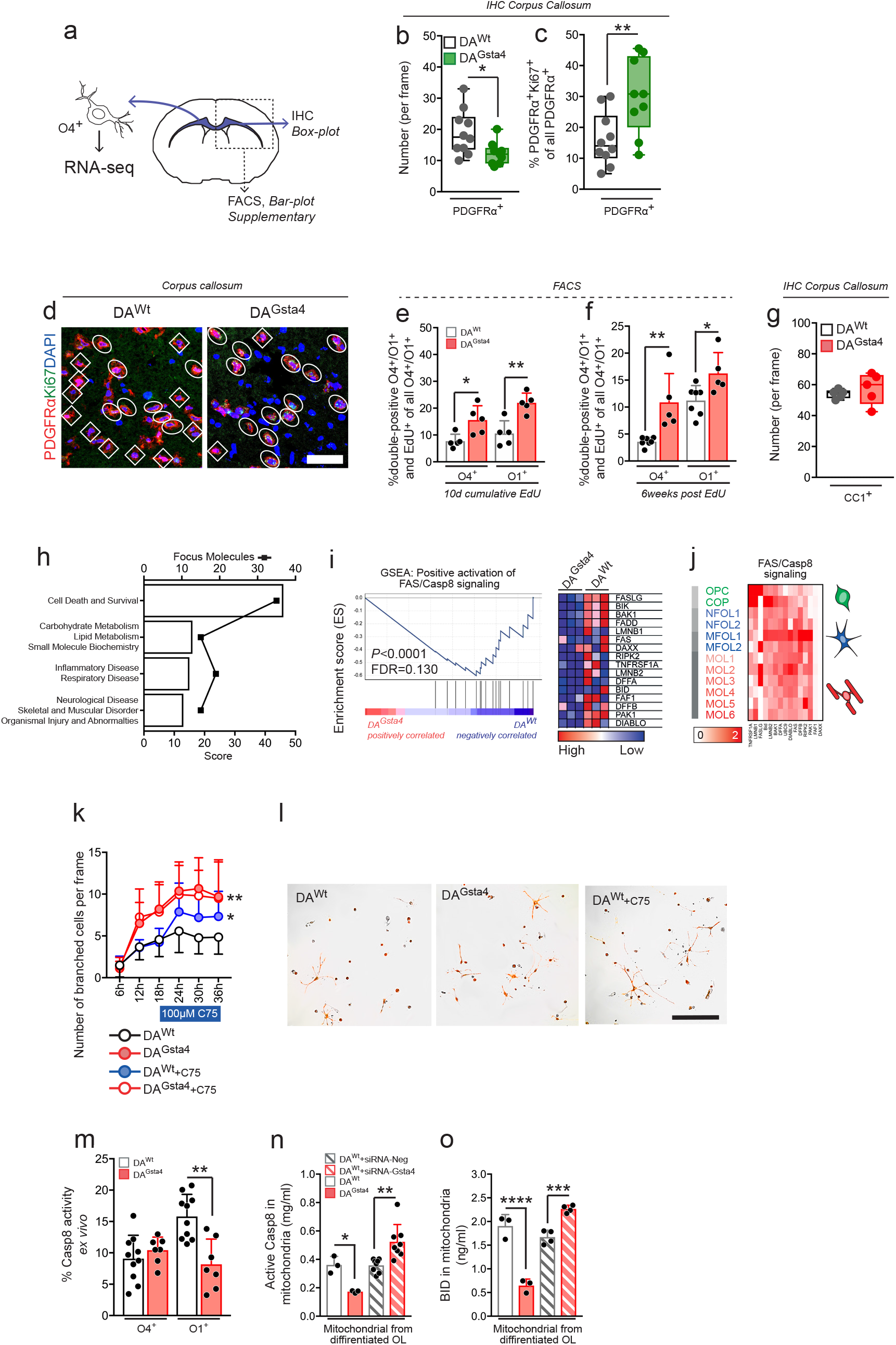
Gsta4 over-expression promotes more efficient OL differentiation via Fas/Casp8/Bid. (a) Summary of experimental set-up. (b) Total number of PDGFRα^+^ OPCs per frame in CC of adult rats (n=10+9). (c) Percentage of double-positive PDGFRα^+^Ki67^+^OPCs of all PDGFRα^+^ OPCs in CC. (d) Representative images illustrating PDGFRα^+^ *(diamond)* and PDGFRα^+^Ki67^+^ *(circle)*, scale-bar 20μ. (e) Percentage of double-positive O4^+^EdU^+^ of all O4^+^ and O1^+^EdU^+^ of all O1^+^ after 10 days of EdU administration (n=5) analyzed with flow cytometry. (f) Percentage of double-positive O4^+^EdU^+^ of all O4^+^ and O1^+^EdU^+^ of all O1^+^ six weeks following EdU administration (n=7+5). (g) Total number of CC1^+^ OLs per frame in CC (n=5). (h) Top IPA Network from RNA-seq on O4^+^ CC OLs (n=5). (i) GSEA and heat map of transcripts promoting Fas/Casp8 signaling from RNA-seq on O4^+^ CC OLs. (j) Heat map based on transcript levels during OL maturation^20^. (k) Number of branched DA^Wt^ and DA^Gsta4^ OLs per image frame captured with live imaging during 36h of differentiation. The Fas inhibitor C75 was added after 18h. (l) representative images after 24h. Overlay with contrast and CellROX staining *(red)*. (m) Percentage of *ex vivo* Casp8 activity in O4^+^ and O1^+^ OLs from DA^Wt^ (n=10) and DA^Gsta4^ (n=7). (n) Quantification of active Casp8 derived from mitochondria of DA^Wt^ and DA^Gsta4^ OLs differentiated for 48h, without (n=3) or with (n=8) siRNA. (o) Quantification of Bid derived from mitochondria DA^Wt^ and DA^Gsta4^ OLs differentiated for 48h, without (n=3) or with (n=4) siRNA. Graphs show representative results from two (e, f, k-o) or one (b-d, g) independent experiments. All graph shows mean and S.D., graph (b-d) shows box and whiskers indicating values outside 5-95 percentile. All statistical analyses were performed with one-way ANOVA apart from (k), analyzed with two-way ANOVA. \**P*<0.05, \*\**P*<0.01, \*\*\**P*<0.001, \*\*\*\**P*<0.0001.

To identify pathways that might contribute to the observed difference in the transition through intermediate OL states, O4^+^ OLs were sorted out from micro-dissected adult DA^Wt^ (n=5) and DA^Gsta4^ (n=5) CC followed by RNA-seq (Fig.2a). Ingenuity Pathway Analysis (IPA) Network suggested top network to be involved in cell survival and death (*Mycn, Egfr, Fas*), carbohydrate and lipid metabolism (*Bicc1, Pisd, Slc25a14*) (Fig.2h, Supplementary Table 1). GSEA on the RNA from the sorted CC O4^+^ OLs also revealed lower expression of positive regulators of Fas and Caspase-8 (Casp8) signaling in DA^Gsta4^ (Fig.2i). A heat map summarizing the levels of transcripts in (Fig.2j) along the OL lineage also showed that transcript levels were higher in early myelin forming OLs (MFOL1) (Fig.2j), suggesting a role in the early stages of differentiation/myelination. To address whether Fas could be beneficial for OL differentiation, the Fas antagonist C75 was added to DA^Wt^ and DA^Gsta4^ OPC cultures during differentiation analyzed with live imaging. DA^Gsta4^ OLs had more branches from the soma as compared to DA^Wt^. However, the DA^Wt^ branching was significantly increased upon addition of C75 (Fig.2k, l).

When assessing intracellular mechanisms downstream of Fas by flow cytometry, Casp8 activity *ex vivo* was indeed lower in O1^+^ DA^Gsta4^ as compared to DA^Wt^ (Fig.2m). No differences were observed in O4^+^ OLs. Casp8 can activate either BH3 interacting-domain death agonist (Bid) or Caspase-3 (Casp3) via catalytic cleavage^57^. Upon cleavage of Bid, both Bid and active Casp8 are associated with the mitochondria outer membrane^57^. We thus quantified the levels of active Casp8 and Bid in isolated mitochondria from differentiated OLs. We observed a decrease in active Casp8 and mitochondrial levels of Bid in DA^Gsta4^ (Fig.2n, o), in accordance to the *ex vivo* findings of reduced Casp8 activity in DA^Gsta4^ OLs (Fig.2m). In addition, knock-down of Gsta4 in differentiated OLs had the opposite effect, with an increase in mitochondrial levels of Casp8 and Bid. Thus, Gsta4 reduces Fas/Casp8 signaling in differentiating OLs, likely to contribute to a more efficient differentiation.

### Gsta4 regulates mitochondrial 4-HNE load in OLs

4-HNE is the sole known substrate for Gsta4 enzymatic activity^58^. In addition, Gsta4 facilitates the extracellular transportation of 4-HNE via conjugation to glutathione whereas additional scavenging enzymes show little overlap with Gsta4-function. Intracellular 4-HNE could thus be considered as a rate limiting molecule on several cellular processes, interfering with normal function, causing protein aggregation and affecting protein degradation^43,44^. Thus, we investigated whether Gsta4 over-expression could also lead to a decrease in intracellular 4-HNE load, resulting in fewer targets being covalently modified by 4-HNE. Staining for 4-HNE in CC revealed the DA^Gsta4^ to have a reduced load of 4-HNE compared to DA^Wt^ (Fig.3a, Fig.S2a). To identify transcripts potentially being directly affected by Gsta4/4-HNE we analyzed the RNA-seq data of CC derived O4^+^ OLs and used the following criteria to identify targets: significant differential expression (*P*=<0.001) between DA^Wt^ and DA^Gsta4^ (Supplementary Table 2) and, the potential of the corresponding peptide to be modified by 4-HNE, proven by high performance liquid chromatography screening^59,60^ (Supplementary Table 3). There were a 10-fold increase in transcripts being upregulated in DA^Gsta4^ OLs compared to DA^Wt^ in naïve state (Fig.3b *left*), however there was no difference in the proportion of transcripts of potentially 4-HNE modified peptides (Fig.3b *green*). Transcripts that were elevated in DA^Gsta4^ included *Ahcy* and *Mat2a*, involved in processes of small carbon units during oxidative stress (Fig.3b *right*). *Fas* and *Bid* involved in Casp8-induced apoptosis was downregulated in DA^Gsta4^ in relation to DA^Wt^, as previously observed (Fig.2h).

**Fig. 3.**
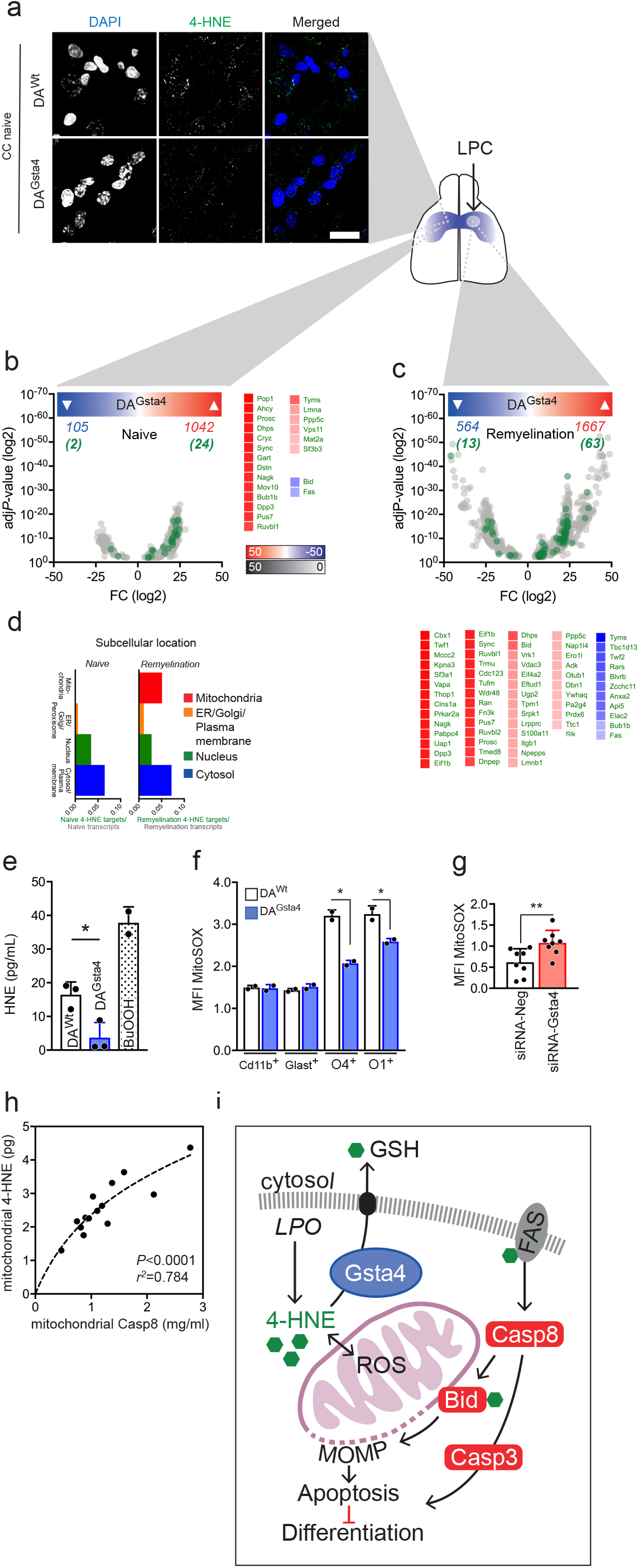
Gsta4 regulates mitochondrial stress and intracellular 4-HNE load in OLs. (a) Staining for 4-HNE in CC of naïve adult DA^Wt^ and DA^Gsta4^ (n=3), scale-bar 10μ. (b) Significant transcripts (*grey*) from RNA-seq of CC derived O4^+^ OLs in naïve conditions. Transcripts corresponding to proteins potentially modified by 4-HNE (*green*)^59,60^. Heat-map indicates up (*red*) and down (*blue*) transcripts. (c) Significant transcripts (*grey*) from RNA-seq of CC derived O4^+^ OLs during remyelination. (d) Subcellular distribution of difference in *green* versus *grey* transcripts in naïve conditions and during remyelination. (e) Mitochondrial 4-HNE load from differentiated DA^Wt^ and DA^Gsta4^ OLs in culture and BuOOH, inducing LPO, as a positive control (n=3). (f) Flow cytometric assessment of mitochondrial integrity with MitoSOX *ex vivo* in glial cells and OLs from DA^Wt^ and DA^Gsta4^ (n=2). (g) Flow cytometric assessment of mitochondrial integrity with MitoSOX in differentiated OLs following application of Gsta4 knockdown with siRNA-Gsta4 (n=8). (h) Correlation between mitochondrial 4-HNE load and levels of active Casp8 associated to the mitochondria (n=16). (i) Schematic illustration of Gsta4-mediated 4-HNE transport and possible binding sites for 4-HNE along the Fas/Casp8 pathway. Graphs show representative results from two (f, h), three (e, g) independent experiments. Data in (e-g) are shown as mean and S.D‥ All statistical analyses were performed using two-tailed Student’s *t* test apart from (h) analyzed with two-tailed Pearson’s *r* test‥ \**P*<0.05, \*\**P*<0.01.

We also assessed Gsta4/4-HNE cross talk in the lysolechitin (LPC) model of induced demyelination/remyelination (Fig.4a). Specifically, we applied stereotactic administrations of 0.1% LPC into the CC and sorted O4^+^ OLs during the remyelination phase. Transcripts including *Mccc2* and *Vdac3* involved in mitochondrial gene expression, were differentially regulated between DA^Wt^ and DA^Gsta4^ (Fig.3c *down*). The subcellular distributions of transcripts of potentially 4-HNE modified proteins (Fig.3b, c, *green*) suggested strain differences in transcript coding for proteins localized to close to the mitochondria during remyelination compared to naïve condition (Fig.3d, Fig.S2d).

**Fig. 4.**
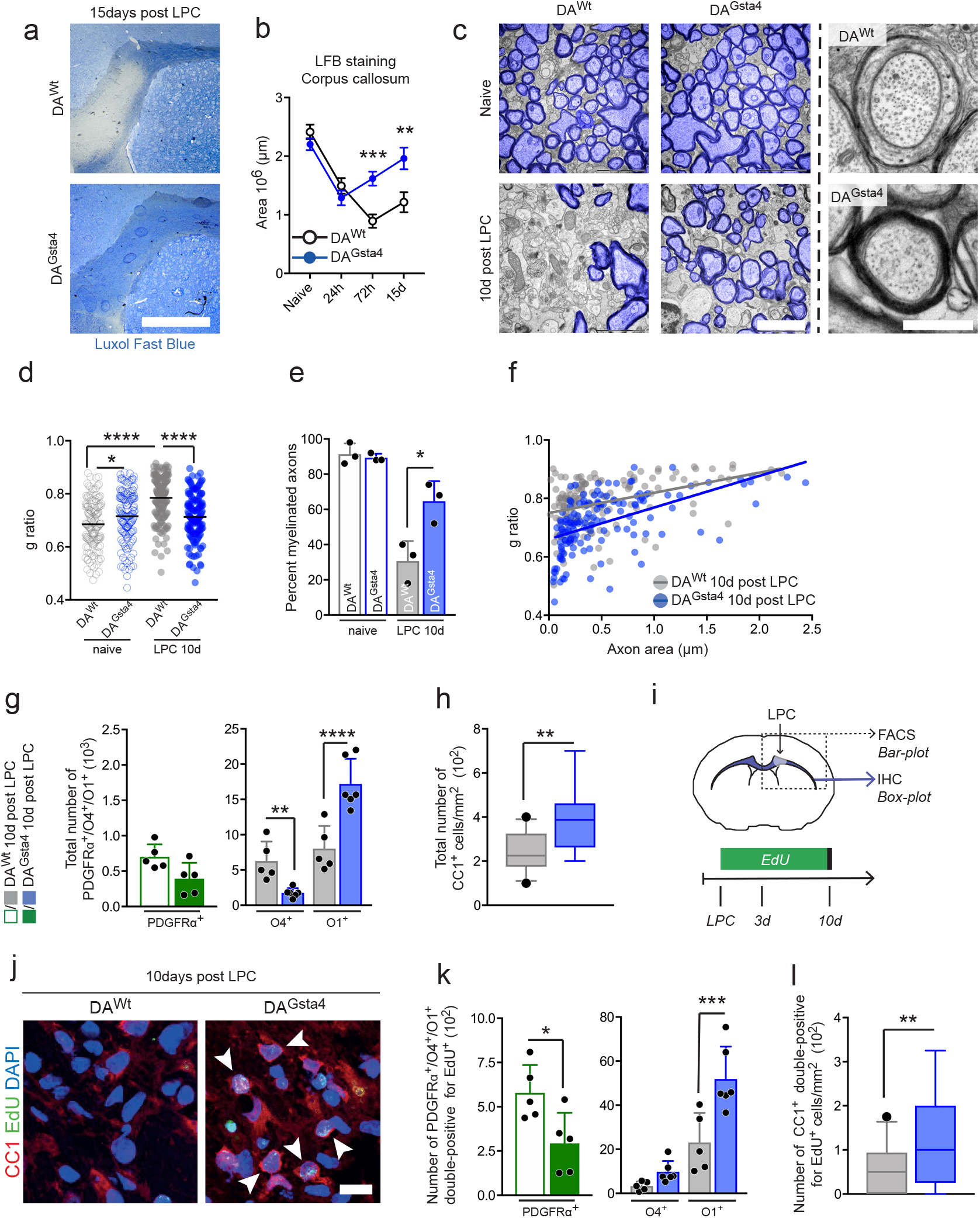
Gsta4 stimulates CC axonal remyelination through more efficient OL differentiation. (a) CC sections stained with LFB for evaluation of demyelination, scale-bar 1mm. (b) LFB staining following administration of 0.1% LPC (naïve/24h/72h; n=6, 15d; n=5). (c) Representative electron microscopy (TEM) images of CC, myelinated axons colored in blue, scale-bar 2μm and 400nm. (d) *G*-ratio in naïve DA^Wt^ and DA^Gsta4^ and during remyelination (n=3). (e) Percentage of myelinated axons in naïve DA^Wt^ and DA^Gsta4^ and during remyelination (n=3). (f) *G*-ratio plotted against axon area following LPC in DA^Wt^ and DA^Gsta4^ (n=3). (g) Flow cytometric analysis of total number of PDGFRα^+^ OPCs *(green)* (n=5) and O1^+^ and O1^+^ OLs *(grey/blue)* (n=5+6). (h) Immunohistochemically validation of CC1^+^ OLs in CC (n=5). (i) Illustration of experimental set-up. (j) Representative CC images ten days following LPC, scale-bar 10μm. (k) Number of PDGFRα^+^, O4^+^, or O1^+^ double-positive for EdU^+^ ten days after continuous EdU administration. (l) Number of CC1^+^ double-positive for EdU^+^ ten days after EdU administration. All graphs show mean and S.D., apart from (h, l) showing box and whiskers, indicating values outside 5-95 percentile. All statistical analyses were performed with one-way ANOVA, apart from (h, l) performed with Students’ two-tailed unpaired *t* test. \**P*<0.05,\*\**P* <0.01,\*\*\**P*<0.001,\*\*\*\**P*<0.0001

As mitochondrial stress and sequent dysfunction is a typical outcome following high intracellular 4-HNE load^33^, we quantified mitochondrial 4-HNE load by isolating mitochondria from OLs differentiated for 48h. DA^Gsta4^ OLs showed reduced mitochondrial 4-HNE load compared to DA^Wt^ and the positive control BuOOH (Fig.3e, Fig.S2b, c), a potent inducer of LPO generating 4-HNE. Mitochondrial stress was measured with MitoSOX *ex vivo* in O4^+^ and O1^+^ OLs and other glial cells (microglia and astrocytes). OLs had higher MitoSOX signal compared to glial cells but DA^Gsta4^ OLs had reduced MitoSOX signal compared to DA^Wt^ (Fig.3f). The signal was then increased upon addition of siRNA-Gsta4 *in vitro* (Fig.3g). Further, mitochondrial 4-HNE load showed a positive correlation with activated Casp8, showing that the beneficial effects of Gsta4 on OL apoptosis involves mitochondrial 4-HNE load (Fig.3h). Taken together our data suggest that Gsta4 over-expression could work as a model for lowering of mitochondrial 4-HNE load and that it could have effects on a wide array of O4^+^ OL transcripts, both during naïve state and during remyelination (Fig.3i, Fig.S2e).

### Gsta4 overexpression facilitates remyelination of CC axons through enhanced OL differentiation

To address if Gsta4 contribute to repair and remyelination in a context of demyelination, we analyzed DA^Gsta4^ rats upon induction of lysolechitin (LPC) mediated demyelination/ remyelination (Fig.4a, Fig.S3a). While Luxol fast blue (LFB) staining initially showed a similar degree of demyelination between the two strains, examinations at later time points revealed a more efficient remyelination in DA^Gsta4^ compared to DA^Wt^ (Fig.4b). The difference between strains was evident already at 72h and persisted until 15 days post injection, a time point when DA^Gsta4^ animals displayed minimal visible demyelinating damage, whereas such damage was still readily evident in the DA^Wt^ strain. No difference in CC size was evident between DA^Wt^ and DA^Gsta4^ (Fig.S3b). Since incomplete scavenging of myelin debris is known to limit OPC differentiation and remyelination, the possible contribution of Gsta4 overexpression or absence during microglia phagocytosis was assessed. pHrodo-labeled myelin was added to primary microglia cultures but no difference in phagocytosis at any time point between DA^Wt^ and DA^Gsta4^ or siRNA^Neg^ and siRNA^Gsta4^ was observed (Fig.S1c, Fig.S3c). In line, transcription of pro-inflammatory hallmark genes did not vary in bulk CC tissue at any time point following LPC exposure (Fig.S3d). This suggests that Gsta4 over-expression had no or limited effect on inflammatory cascades in the LPC model. To evaluate remyelination of axons, naïve CC and CC 10 days following LPC injection were evaluated using transmission electron microscopy (TEM) (Fig.4c, Fig.S3e). The *G*-ratio describes the diameter of the transected axon in relation to the outer diameter of the surrounding myelin depicted by TEM. By using a software, randomly selecting axons for analysis from every image frame^61^, DA^Wt^ axons was found to be less myelinated after remyelination compared to DA^Gsta4^ (Fig.4c, d, Fig.S3e). A small but significant difference was also found already between naïve animals, where DA^Gsta4^ axons had slightly thinner myelin. A two-fold increase in the percentage of myelinated axons was recorded in DA^Gsta4^ during remyelination (Fig.4e) and the difference in myelin thickness was primarily evident in thinner axons (Fig.4f). In line with more efficient maturation (Fig.2) and more rapid remyelination in DA^Gsta4^, DA^Gsta4^ also generated a larger number of O1^+^ and CC1^+^ cells compared to DA^Wt^ 10 days after LPC (Fig.4g-h, Fig.S3h). The total number of O4^+^ OLs and PDGFRα^+^ was decreased in DA^Gsta4^ compare to DA^Wt^ following LPC, supporting the theory of a faster remyelinating process in DA^Gsta4^. To validate this, animals were fed EdU in their drinking water for 10 days (Fig.4i, j, Fig.S3f, g) showing a higher number of DA^Gsta4^ OPCs differentiated into myelin producing O1^+^ OLs or CC1^+^ OLs compared to DA^Wt^ (Fig.4j-l).

### Gsta4 reduces demyelinated lesions and ameliorates EAE

To address the implication of Gsta4 overexpression in a model of autoimmune demyelination we induced experimental autoimmune encephalomyelitis (EAE) in DA^Wt^ and DA^Gsta4^ by sub-cutaneous injection of recombinant myelin oligodendrocyte glycoprotein (MOG) in adjuvant. There was no difference in terms of disease incidence, onset or mortality between the groups (Fig.S4a), suggesting that priming of infiltrating leukocytes did not differ between the strains. Animals over-expressing Gsta4, however, displayed significantly improved clinical signs (Fig.5a), reduced disease duration and cumulative disability scores compared to DA^Wt^ (Fig.5b). At the end of experimental follow up, DA^Gsta4^ also showed reduced mitochondrial stress assessed with MitoSOX in O1^+^ cells without any evident differences in mitochondria presence (Fig.5c, d), labeled with MitoTracker. No differences in the numbers of infiltrating leukocytes or activated microglia in the spinal cords between DA^Gsta4^ and DA^Wt^ was observed (Fig.S4d) which supports the notion of Gsta4 playing no or a minor role for regulating adaptive or innate immunity. In order to further substantiate this notion, we performed EAE experiments in bone marrow transplanted DA^Wt^ and DA^Gsta4^ (Fig.S4b, c). Animals were lethally irradiated and transplanted with green fluorescent protein (GFP)-expressing bone marrow from DA^GFP^. DA^Gsta4^ animals transplanted with DA^GFP^ bone marrow (DA^GFP^→DA^Gsta4^) recapitulated the EAE phenotype previously observed in DA^Gsta4^ (Fig.5e). Disease duration and maximal disease scores were significantly lower in DA^GFP^→DA^Gsta4^ compared to DA^GFP^→DA^Wt^ (Fig.5f). In addition, DA^GFP^→DA^Gsta4^ showed lower levels of active Casp8 staining in CC1^+^ spinal cord (Fig.5g, h). In line with our previous findings that restricted Casp8 activity contributes to OL maturation DA^Gsta4^ also displayed smaller area of demyelination and fewer lesions in spinal cord dorsal column (Fig.5i, j). Taken together, this set of experiments suggest that Gsta4 over-expression contributes to a lowering of active Casp8 and mitochondrial stress which results in fewer demyelinated lesions or increased remyelination, and improved clinical symptoms.

**Fig. 5.**
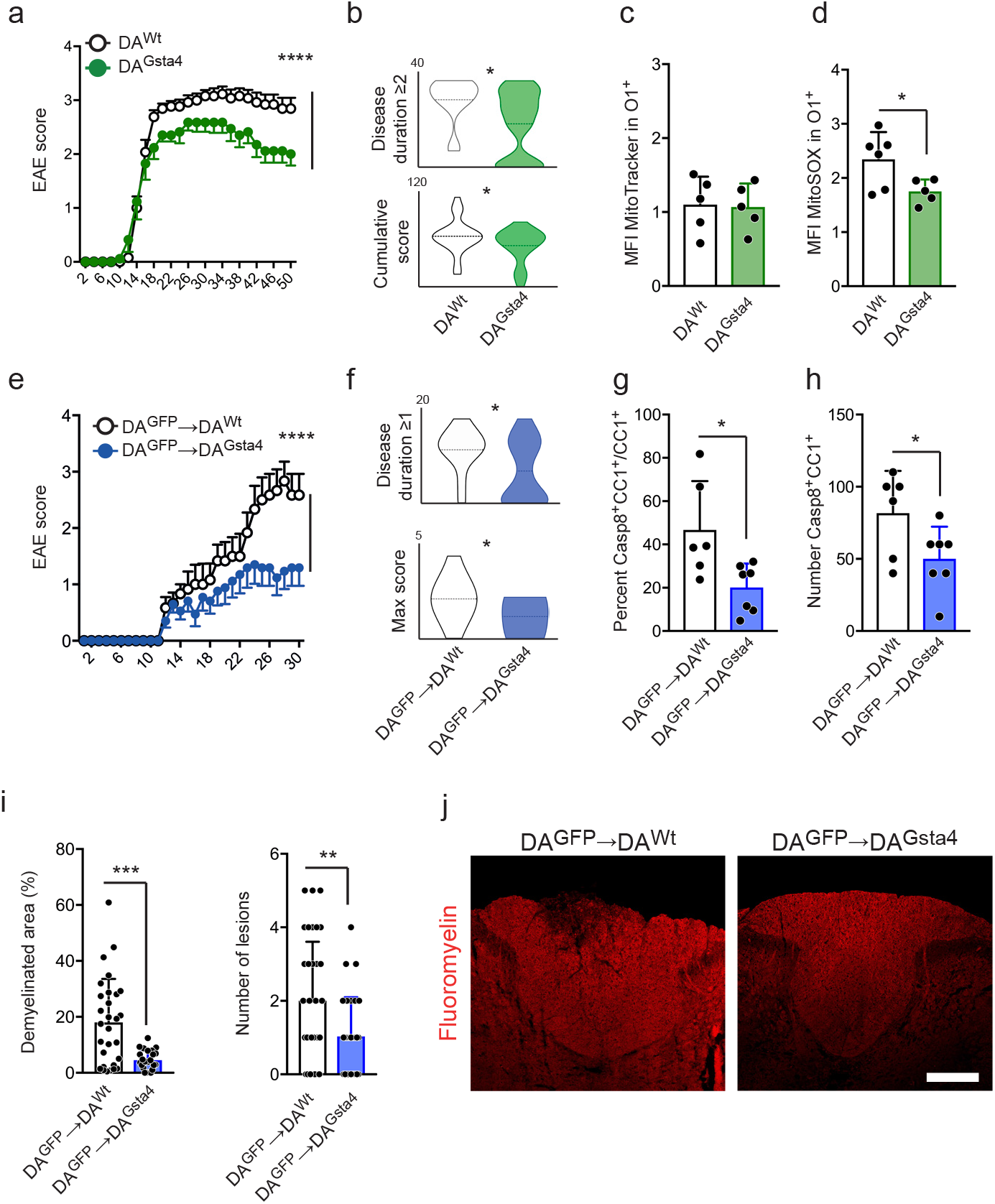
Gsta4 in CNS ameliorates EAE and reduce demyelinating lesions. (a) EAE was induced with subcutaneous injection of MOG in adjuvant (n=26+17). (b) Disease duration and cumulative score based on (a). (c, d) Levels of MitoTracker and MitoSOX in O1^+^ OLs from spinal cord at day 50 from (a). EAE in lethally irradiated DA^Wt^ (n=12) and DA^Gsta4^ (n=16) transplanted with DA^GFP^ bone marrow. (f) Disease course and max score based on (e). (g, h) Percentage of Casp8^+^CC1^+^ of all CC1^+^ (n=3) and number of Casp8^+^CC1^+^ OLs in spinalcord (n=3) at day 30 based on representative animals in (e). (i) Demyelinated area of total area of dorsal column and number of dorsal column lesions (n=3) at day 30 based on representative animals in (e). (j) Representative images, scale-bar 100μm. All data are shown as mean and S.D. apart from (b, f) showing violin plots and mean. Two-way ANOVA test was used for (a, e) to analyze differences between groups over-time, graphs are showing mean and S.E.M. Remaining statistical analyses were performed using Student’s two-tailed unpaired *t* test, apart from disease duration and number of lesions analyzed with Mann Whitney test. \**P*<0.05,\*\**P* <0.01,\*\*\**P*<0.001,\*\*\*\**P*<0.0001

## Discussion

Some progress has been made in the identification of signaling pathways that could be of substantial relevance for OL maturation and clinical applications. These include the Wnt pathway, interfering with remyelination in the mammalian CNS ^62^ and retinoid acid receptor RXR-γ signaling that enhances OPC differentiation and remyelination ^63^. A large scale high-throughput screening of compounds with remyelinating effects identified a cluster of anti-muscarinic compounds, including CF, that enhanced OPC differentiation and remyelination *in vitro* ^54^. CF, a first generation anti-muscarinic compound and recently identified as a promising candidate for remyelinating therapy in the clinical settings ^54–56^. We show CF and DMF to promote OL differentiation in a Gsta4-dependent manner, and that this involves Nrf2. The anti-muscarinic compound contains fumarate and thus makes it partly the same substance as DMF, where both compounds have the same fumarate molecule. This was the notion for addressing the overlapping features between CF and DMF to activate Nrf2 in OLs. We thus speculate that the fumarate molecule of CF is likely the cause of Nrf2 activation, but to what extent it is a combined effect of clemastine and fumarate needs to be further investigated. However significant decline in OL differentiation following DMF and CF application in the absence of Gsta4 illustrates the importance of scavenging enzymes and cellular protection during differentiation.

Despite detailed knowledge of the molecular heterogeneity between juvenile and adult OL maturation ^64^, there are still large gaps in our understanding of what stops OLs from remyelinating the damaged adult CNS. We herein suggest newly formed/myelin forming OLs to represent a particularly sensitive state during adult OL differentiation and consequently be a rate limiting state during maturation that can benefit from elevated Gsta4 levels (Fig.2i, j, Fig.3e). Our approach to identify pathways affected by Gsta4 over-expression and sequent reduction of mitochondrial 4-HNE load was to merge two independent databases on 4-HNE modified proteins together with transcription in OLs.

The transcripts for *Fas* and *Bid* were the only significantly downregulated in DA^Gsta4^ OLs compared to DA^Wt^ (Fig.3b). Both Fas and Bid protein have been described to be 4-HNE modified. Fas can be modified by 4-HNE at three different sites, including one modification within its active site (Fig.S2e). We confirmed that lowering of Fas (activity) was beneficial to OLs since addition of Fas inhibitor C75 lead to an increase in the number of branched OLs in DA^Wt^ but not DA^Gsta4^ OLs. The involvement of this pathway was further illustrated by reduced activity of Casp8 in DA^Gsta4^ OLs *ex vivo*. Both mitochondrial Casp8 and Bid were negatively correlated to Gsta4 expression, further proving the mitochondria as a relevant location for the intersection between 4-HNE and additional pathways, including the Fas/Casp8/Bid (Fig3e, h). The decline in 4-HNE load could be of importance for certain maturation checkpoints during OL differentiation and enable more efficient intracellular processes.

Interestingly Gsta4 also affected OPC proliferation both in neonatal-derived cultures and adult rats. However, the Gsta4-mediated differentiation appeared superior the proliferation able to reduce the number of OPCs despite a higher proportional proliferation compared to DA^Wt^. Collectively, our findings suggest that Gsta4 and physiological levels of 4-HNE regulate the differentiation of OLs and that this has clinically relevant effects in *in vivo* models of inflammation-or toxin-induced demyelination. Furthermore, we propose that the myelination-promoting effects of both DMF and CF involve the Gsta4/4-HNE pathway and that the Gsta4 is essential for beneficial outcome when applying these compounds. Further studies are needed to explore the role of Gsta4 in a clinical context and if pathways identified herein can also be of importance in man and addressed as possible therapeutic targets.

Tracking of maturing OLs by EdU incorporation after focal demyelination supports the notion that DA^Gsta4^ OLs more rapidly advance through intermediate states of differentiation to more rapidly make up a higher number of myelinating cells to enclose damaged CC axons (Fig.4). The CC of DA^Gsta4^ is nearly completely restored 15days following focal demyelination whereas DA^Wt^ still have tangible lesions. This is also in line with TEM analysis, where we observed that DA^Gsta4^ have the double amount of myelinated axons 10 days following demyelination. This is despite a similar initial demyelination in both strains (Fig.4b). Interestingly, DA^Gsta4^ axons in naïve adult animals appeared to have a slightly thinner myelin sheet. The OPC pool is also reduced in DA^Gsta4^, and an explanation for this and for the thinner myelin sheet could be that upon release of the 4-HNE brake cellular processes will operate more efficient and a smaller pool of precursor cells or thinner myelin is then sufficient to fulfill the function of metabolic support and axonal transmission ^65^.

Lastly, we observed that Gsta4 over-expression leads to behavioral amelioration upon induction of MS-like disease (EAE). Gsta4 did not affect time of onset or incidence but improved disease duration and cumulative and max disability scores, also upon depletion of Gsta4 over-expression from the immune system. This could further be linked to mitochondria dysfunction and Casp8 which both were reduced in DA^Gsta4^ OLs.

Facilitating and increasing remyelination has been a longstanding therapeutic goal in chronic demyelinating diseases such as MS ^66^. Until very recently, successful examples especially in the clinical setting has been all but lacking. Our study places Gsta4 as a key regulator of OL differentiation and remyelination in the context of cellular stress, ageing and demyelination, as well as further elucidates the mode of action of two promising pro-myelinating agents, DMF and CF.

## Methods

### Study Design

Our research object was to investigate the possible beneficial role of Gsta4, activated by DMF and Clem-F, in remyelination and *in vivo* OL differentiation in MS disease-like models. The DA^Gsta4^ rat was thus designed and used to over-express Gsta4 both in primary cultures and in *in vivo* models in which siRNA towards Gsta4 was used as a knock-down tool. All primary OL cultures were established from neonatal rats (3-7days) whereas adult animals at an age of 8-10 weeks were used for bone marrow transplantation and later EAE after an additional 8-10 weeks. For remaining experiments adult animals at an age of 10-12 weeks were used. Throughout this study littermate controls were used and all caged contained animals with different genotypes/treatment. OL differentiation was evaluated in the corpus callosum using PDGFRα^+^ to label OPC and CC1^+^ to label post-OPCs. Upon handling and evaluation with IHC and LFB the investigator was unaware of the genotype of the animals/sections.

### Animals and genotyping

The rat DA^Gsta4^ strain over-expressing *Gsta4* under a CAG-promotor was purchased from Taconic Bioscience. DA^GFP^ animals were kindly provided by Holger Reichardt’s laboratory and have been described previously (Danielyan L, et al. 2009). Animals were genotyped using PCR and vector-specific primers (F: ATCCACTTTGCCTTTCGCGCC, R: TTTCAAACACTGGGAAGTAACG), and primers towards Cd79b (F: GACTCTGTGGCTGCTCATCC, R: TTCAGCAAGAGCTGGGGAC) as wild-type allele control. Animals were bred in the animal facility at Karolinska University Hospital (Stockholm, Sweden) in a pathogen free and climate-controlled environment with regulated 12h light/dark cycles. All experiments were approved and performed in accordance with Swedish National Board of Laboratory Animals and the European Community Council Directive (86/609/EEC) under the permits N275-15 and N244-13.

### Bone marrow transplantation and experimental autoimmune encephalomyelitis (EAE)

DA^GFP^ male animals were sacrificed with carbon dioxide and femur was removed and rinsed with cold PBS in order to harvest hematopoietic stem cells (HSC). These were washed, passed through a 40μm strainer and erythrocytes were lysed using ACK buffer (Thermo Fisher Scientific, Waltham, MA) following the manufacturer’s protocol. At an age of 8-10weeks, male BM recipients were subjected to lethal irradiation with 2×5Gy and subsequently given 10×10^6^cells/in 300μL PBS i.v. Successful transfer of the bone marrow graft was assessed by analyzing GFP^+^ cells in the peripheral blood 8 weeks after transplantation using flow cytometry. EAE in bone marrow transplanted male animals was induced in anesthetized 8-10 week-old animals after irradiation by subcutaneous injection at the tail base of 10μg of recombinant rat MOG together with incomplete Freud’s adjuvant 1:1 (Sigma, St Louis, MO). EAE was induced in 10-12weeks old female animals using the same protocol using 10μg MOG. The animals were monitored daily from onset of disease and scored according to the following scheme: 0=no clinical score, 1=reduced tail tonus, 2= hind leg paraparesis or hemiparesis, 3=hind leg paralysis or hemiparalysis, 4=tetraplegia or moribund, 5= death. Rats were sacrificed if they showed severe balance disturbance or abnormal behavior, >25% weight loss or prolonged tetraplegia. Removed animals were throughout scored as 4. EAE experiments were repeated twice (EAE1 n=17+26; EAE2 n=16+18) and twice in bone marrow transplanted animals (BM-EAE1 n=12+16; BM EAE2 n=10+14) with comparable and significant results. For oral gavage with DMF, animals were given vehicle or DMF (150 mg/kg) in 0.1% methyl cellulose via oral gavage in a total volume of 1 mL (Sigma, St. Louis, MO). Brains were collected an bulk mRNA isolated as described below 5 h after administration.

### Lysolecithin injections

Male rats animals 10-12 weeks were anaesthetized with isoflurane and subjected to a stereotactic injection into the corpus callosum (AP: 1.0mm; L:1.2; DV: 2.2) with 2.5μL 0.1% lysolecithin (Sigma, St Louis, MO) in PBS over 5 min using a Hamilton syringe 1710RN (Sigma, St Louis, MO)^67,68^. Animals were kept for 24h, 72h, 10d or 15d and evaluated with flow cytometry, qPCR, TEM or immunohistochemistry. For *in vivo* differentiation, 5-ethynyl-2’-deoxyuridine (EdU; Thermo Fisher Scientific, Waltham, MA) was either injected i.p. 25mg/kg, added to the drinking water (0.2mg/mL) for a period of 10d following LPC injection or in naïve animals for the same time period. Injected animals were kept for 6weeks.

### Primary OPC cell cultures

For rat primary OPC cell culture preparation brains were collected from neonatal rats (3-7d). The tissue was incubated in 1mL Accutase and 4μL DNase/mL (Sigma, St Louis, MO) at +37°C and passed through a fire–polished Pasteur pipette every 10^th^ min, this being repeated three times with decreasing pipette guage. The homogenate was directly labeled and isolated using magnetic anti-A2B5 beads (130-093-388) (Milteny Biotec, Bergisch Gladbach, Germany) instead of Percoll layering as for flow cytometry. Throughout the study, culture ware was pre-coated with Poly-L-lysine (Sigma, St Louis, MO) at +37°C for >2h, rinsed with distilled water twice and incubated with Fibronectin (Sigma, St Louis, MO) at +37°C for >1h. Cells were cultured in 200μL Neurobrew (Milteny Biotec, Bergisch Gladbach, Germany), 100μL N2-supplement (Thermo Fisher Scientific, Waltham, MA), 2μL bFGF (PeproTech, Princeton, NJ) and 2μL PDGF-BB (R&D Systems, Minneapolis, MA) per 10mL of DMEM/F-12 media (Thermo Fisher Scientific, Waltham, MA). For differentiation bFGF and PDGF-BB was removed from the media. Cells were proliferated for 5-7d at +37°C and 5% CO_2_ in a humidified incubator, harvested with Accutase and re-seeded; 50 000cells/well for 96-well plate (Nunc, Rochester, NY) or 75-100 000cells/well for staining in chambers (177402PK) (Nunc, Rochester, NY).

### siRNA targeted knockdown

For siRNA knockdown samples were transfected with 1pmol siRNA together with 0.3μL Lipofectamine (Thermo Fisher Scientific, Waltham, MA) per 200μL and incubated for 24h in media. siRNA sequences were design using Thermo Fisher online software, siRNA-Gsta4 (#15) (used throughout): sense; GCAUUUAAGACAAGAAUCAtt; antisense; UGAUUCUUGUCUUAAAUGCct, siRNA-Gsta4 (#22): sense; GGAUGGAUGCCUGCUUUUUtt, antisense; AAAAAGCAGGCAUCCAUCCtt, siRNA-Neg (#4404021) (Thermo Fisher Scientific, Waltham, MA).

### OL stimulation with DMF, Clem-F, Clem-H and C75

DMF (PubChem ID: 637568) and Clem-F (PubChem ID: 5281069) (Sigma, St Louis, MO) and Clem-H (PubChem ID: 66847663) (BOC Science, Shirley, NY) were dissolved in DMSO and added to differentiation media at a final concentration of 10μM 24h. an equal amount of DMSO was added to control samples. OL differentiation was quantified with RT-qPCR and anti-Plp staining, described below. The Fas inhibitor C75 (Sigma, St Louis, MO) was dissolved in DMSO and added to differentiation media at a final concentration of 100μM for 18h following 18h of differentiation. An equal amount of DMSO was added to control samples. When switching to differentiation media OLs were labeled with CellROX Deep Red (Thermo Fisher Scientific, Waltham, MA) at a final concentration or 1000nM. OL differentiation was quantified with live cell imaging using IncuCyte™ ZOOM (Essens Bioscience, Göttingen, Germany). Deep red signal and contrast were acquired every second hour for 36h. The number of branched cells was used as a measurement for OLs differentiated cells.

### Immunocytochemistry

For immunocytochemistry cells were washed and fixed in 100μL 4% PFA at +4°C for 10min, washed with PBS and incubated with 0.1% Triton X-100 for 5 min at +4°C. For Plp-staining cells were washed and stained with anti-Plp 1:500 (NB100-1608) (Novous Biologicals, Littleton, CO) overnight followed by anti-chicken-Cy3 (Thermo Fisher Scientific, Waltham, MA) and co-stained with CellMask and Hoechst (Thermo Fisher Scientific, Waltham, MA) for 5min at +4°C. Co-localization was also performed according to the manufacturer’s protocols using Puromycin (Sigma, St Louis, MO) as a negative control and anti-HNE (ab48506) (Abcam, Cambridge, GB) (1:50). Confocal images were for above described applications were acquired with a DMI6000 microscope (Leica Biosystems, Wetzlar, Germany) and analyzed using ImageJ default plugins for co-localization and staining intensity.

### OPC proliferation

Primary cultures of OPC derived from neonatal rats were expanded and reseeded 500 000cells/well in a 48-well plate. Settled cells were cultured with 10μM EdU for 72h in media containing Neurobrew, N2 and PDGF-BB as described above. In addition, media was supplemented with full (1/1), half (1/2) of absent (0) of bFGF concentration referred to the concentration described above. Cells were harvested with Accutase, labeled with anti-PDGFRα (Milteny Biotec, Bergisch, Germany). Dead cells were excluded using near IR Live/Dead (Thermo Fisher Scientific, Waltham, MA). Samples were analyzed with a 3-laser Beckman Coulter Gallios using Kaluza Software (Beckman Coulter, Brea, CA).

### Microglia phagocytosis

For primary microglial cultures brains were collected from rats 10-12 weeks as described above. The cell pellet obtained after Percoll layering was plated in non-precoated culture ware for adherent cells and expanded for 4 days in DMEM/F-12 media with 10% FCS (Sigma, St Louis, MO). Microglia were isolated using anti-Cd11b beads (130-049-601) (Milteny Biotec, Bergisch Gladbach, Germany) according to the manufactures’ exact instructions. pHrodo labeled rat-myelin was kindly provided by Rasmus Berglund, Karolinska Institute, and was added to the media for 3h and 6h followed by flow cytometric analysis as described above using anti-CD11b (WT5. ROU) (BD, Franklin Lakes, NJ).

### Flow cytometry and Caspase-8 activity

Animals were sacrificed with carbon dioxide and perfused with PBS, the brain was removed and minced using a sterile scalpel. Tissues were processed as described for primary OPC cultures. Following enzymatic digestion, the homogenate was filtered through a 70μm strainer and myelin was removed by a 37% Percoll layer spun at 800 x g for 10 min at +10°C without any brake. The pellet was re-suspended to a single cell suspension and stained for EdU Click-IT (Thermo Fisher Scientific, Waltham, MA) according to manufacturer’s instructions and then co-stained with anti-PDGFRα (KG215-7) (Creative Diagnostics, NY), anti-O4 (130-118-978) (Milteny Biotec, Bergisch, Germany), anti-O1 (FAB1327V) (Novous Biologicals, Littleton, CO) and anti-CD45 (OX-1 (ROU)) (BD, Franklin Lakes, NJ). Dead cells were excluded using near IR Live/Dead (Thermo Fisher Scientific, Waltham, MA). Caspese-8 activity was measured by *ex vivo* incubation of single cell suspension, prior to Ab-staining described above. The single cell suspension was incubated for 1h at 37°C in 300uL DMEM/F12 media, supplemented with N2 and Neurobrew. FAM-LETD-FMK Caspase 8 reagent (Thermo Fisher Scientific, Waltham, MA) was added at the concentration described by the manufacturer and stained. Samples were analyzed using a 3-laser Beckman Coulter Gallios using Kaluza Software (Beckman Coulter, Brea, CA).

### Cell sorting, microarray and RNA-seq

Animals were sacrificed and perfused as described above, corpus callosum was dissected and prepared as samples for flow cytometry. The singe cell suspension was labeled with magnetic anti-CD11b (130-049-601) and anti-Glast (130-095-825) beads at a dilution suggested by manufacturer and incubated and separated using LS columns. The negative fraction was stained with anti-O4 beads (130-094-543) and isolated using MS columns. Beads, reagents and columns were provided by Milteny Biotec, Bergisch Gladbach, Germany. RNA was isolated using RNeasy Micro Kit (Qiagen, Venlo, Netherlands) according to the manufacturer’s instructions with minor modifications to the protocol. In brief, isolated cells were lyzed in 500μL Qiazol following sorting, 100μL chloroform were added and samples spun at 7000rpm for 15min at +4°C in a MaXtract column (Qiagen, Venlo, Netherlands). Mixed with 500μL 70% EtOH and spun at 10 000rpm for 30s at +4°C in a MiniElute column, washed with 350μL RW1, on-column digestion for 15 min at room temperature, washed with 350μL RW1, 500μL RLT, followed by 500μL 80% EtOH and spun for 2min. RNA was eluted using 10mM Tris pH7.5. Buffer. Next generation sequencing and bioinformatics analysis was performed by the National Genomics Infrastructure (NGI) at the Science for Life Laboratory on the HiSeq 2500 System using the HiSeq Rapid SBS Kit v2 (Illumina, San Diego, CA) generating >13.5 M reads/sample. Reads were mapped to the Rnor_6.0 reference genome using STAR (Dobin et al 2016). Data normalization and analysis of differential gene expression was done using the DESeq2 R-package (Love et al 2014) using a negative binomial test. The false discovery rate-adjusted P-value (FDR) was estimated based on Benjamin-Hochberg correction. For microarray and qPCR analysis of bulk tissue from naïve DA^Gsta4^ (n=4) and DA^Wt^ (n=4) animals, including cortex, hippocampus and corpus callosum was prepared as the sorted samples using a tissue raptor (Qiagen, Venlo, Netherlands) for lyzing in Qiazol. Sub sequent steps were performed similar to RNA extraction form sorted cells described above. Samples were analyzed on a RaGene-2_1.st array (Affymetrix, Santa Clara, CA) by the Array and Analysis facility at Uppsala University. GSEA was performed using http://www.genomespace.org and gene set GO_POSITIVE REGULATION_OF_OLIGODENDROCYTE_DIFFERENTIATION and were calculated by GSEA with weighted enrichment statistics and ratio of classes for the metric as input parameters. Prediction of subcellular locations was generated based on Supplementary Table 2, 3 using www.subcellbarcode.com.

### Histopathological analyses and immunohistochemistry

Animals were sacrificed with carbon dioxide and perfused with PBS followed by 4% formaldehyde and post-fixed in formaldehyde for 24h followed by 3d in 40% sucrose, snap frozen in iso-pentane and kept at −80°C. Sections (12μm) were prepared in a cryostat and kept at −20°C. The EdU staining (Thermo Fisher Scientific, Waltham, MA) was done according to manufacturer’s instructions. For additional staining’s, sections were blocked in 3% normal goat serum (Sigma, St Louis, MO) for 1h, repeatedly washed with PBS and incubated with primary antibodies; anti-PDGFRα (ab5460) (1:100), anti-CC1 (OP80) (1:100) (Millipore, Darmstadt, Germany), anti-Casp8 (1:200) (NB100-56116) (Novous Biologicals, Littleton, CO) or anti-HNE (1:50) (ab48506) (Abcam, Cambridge, GB) overnight. All secondary antibodies were produced in goat and labeled with either Alexa Fluor 488, 594, Cy3 or Cy5 (Thermo Fisher Scientific, Waltham, MA). Following PBS wash secondary Abs were incubated 1:400 for 1h at room temperature. Sections were again washed and mounted with medium containing DAPI (Sigma, St Louis, MO). For LFB (Sigma, St Louis, MO), sections were dried for 30min, hydrated in 95% EtOH for 2min, incubated 2h in LFB at +60°C, washed in dH_2_O, then washed in 95% EtOH, dH_2_O, freshly prepared 0.05% Li_2_CO_3_, 70% EtOH and washed in H_2_O. Sections were analyzed using a Nikon Eclipse E600 microscope and scanned using a Nikon LS-2000 film scanner (Nikon, Minato, Japan). The LFB staining was analyzed using ImageJ software by measuring corpus callosum size and LFB intensity in black/white transposed images.

### Transmission electron microscopy (TEM) and analysis

For TEM evaluation of corpus callosum rats were perfused as described above using 2.5% glutaraldehyde, 1% paraformaldehyde in 0.1M PBS. Rinsed with 0.1PBS and post-fixed in 2% OsO_4_ in 0.1M PBS at +4°C for 2h. Brains were dehydrated in 70%-OH for 30min +4°C, 95%-OH for 30min +4°C, 100%-OH 20min RT, Acetone 2×15min RT, LX-112/Acetone (1:2) 4h RT, LX-112/Acetone 1:1 overnight RT, LX-112/Acetone (2:1), overnight RT, LX-112 overnight RT. Embedding in LX-112 at +60°C. Embedding and sectioning were performed and pictures taken by EMiL, Clinical Research Center, Department of Laboratory Medicine, Karolinska Institute, Huddinge, Sweden. 150 axons from 3 animals per condition were sub-sequentially analyzed randomly and blinded using ImageJ and *G*-ratio free plugin^61^.

### Mitochondria isolation and ELISA

Mitochondria was isolated from primary OL cultures using Mitochondria Isolation Kit for Cultured Cells (#89874) (Thermo Fisher Scientific, Waltham, MA) as per the manufacturers exact instruction. The lysed mitochondria were kept at −80°C until analysis and diluted 1:1 in sample diluent. Protein from whole primary OL cultures were obtain by lysing cells in RIPA buffer (Thermo Fisher Scientific, Waltham, MA) and sub sequential sonication after removal of media and repeated wash with PBS. The lysed cells were kept at −80°C until analysis and diluted 1:1 in sample diluent. α-Tocopherol (50μM) and tert-Butyl hydroperoxide (25μM) (Sigma, St Louis, MO) were used as negative and positive controls and added for 2h to the cultures. The Casp8 ELISA (EKR1606) (Nordic Biosite, Sweden), Bid ELISA (NBP2-69968) (Novous Biologicals, Littleton, CO) and the competitive ELISA for detection of 4-HNE (CSB-EQ027232RA) (Cusabio Technology, Huston, TX) was performed due to the manufacturers exact instruction.

### Quantitative real-time PCR

Total RNA was isolated from tissue or cells using Qiazol and RNeasy mini kit (Qiagen, Venlo, Netherlands) according to manufacturer’s instructions and with 15min on-column DNase digestion (Qiagen, Venlo, Netherlands). cDNA was prepared with reverse transcriptase using iScript kit (BioRad Laboratories, Hercules, CA). Amplifications were conducted using Bio-Rad SYBR green according to manufacturer’s instructions and plates were run in Bio-Rad CFX optical system (BioRad Laboratories, Hercules, CA). Primers were design to work at +60°C and to span an exon junction using online software at http://www.ncbi.nlm.nih.gov. Primer specificity was assessed by determining amplicon size using gel electrophoresis and melt curve analysis of each reaction indicating a single peak. *Bactin* F: CGTGAAAAGATGACCCAGATCA; R: AGAGGCATACAGGGACAACACA, *Hprt*: F: CTCATGGACTGATTATGGACA; R: GCAGGTCAGCAAAGAACTTAT, *Gsta4* F:CAGGAGTCATGGAAGTCAAAC; R: TTCTCATATTGTTCTCTCGTCTC, *Plp1* F: TTGGCGACTACAAGACCACC; R: TGTACACAGGCACAGCAGAG, *Il6* F:AGAAAAGAGTTGTGCAATGG; R:ACAAACTCCAGGTAGAAACG, *Tnf* F:CCGTCCCTCTCATACACTGG; R: GGAACTTCTCCTCCTTGTTGG, *iNos* F: CAACATCAGGTCGGCCATTACT; R: TAGCCAGCGTACCGGATGA, *Mbp* F: ACACACAAGAACTACCCACTACGG; R: GTACGAGGTGTCACAATGTTCTTG. Ireg1 F: CAGCTTTGCTGTTCTTTGCCT; R: AGACGCTCTCCCCTTGTTTG.

### Statistical analysis

General statistical analyses were performed in GraphPad Prism software. Two group comparisons were throughout performed using Student’s two-tailed unpaired *t* test. Two group comparisons with a control group were analyzed using one-way ANOVA. Comparisons in EAE and stimulation with C75 were conducted with two-way ANOVA. *P*<0.05 was throughout considered statistically significant. Analysis of RNA-seq data was performed using the DESeq2 package, which uses Wald testing with Benjamini–Hochberg adjustment for multiple testing.

## Acknowledgement

We thank R Berglund for providing fluorescence myelin and experiment design. We thank the staff at AKM for animal caretaking, especially A Ilveus and A Laurell. We thank K Hultenby and E Idsund Jonsson for their expertise in preparing tissues and taking TEM images at EMiL. We acknowledge support from the Science for Life Laboratory, the National Genomics Infrastructure (NGI) and Affy and Analysis Facility in Uppsala for providing assistance in RNA-seq and microarray sequencing. Work in GCB.’s research group was supported by Swedish Research Council (grant 2015-03558),EuropeanUnion (Horizon 2020 Research and Innovation Programme/EuropeanResearch Council Consolidator Grant EPIScOPE, grant agreement number 681893), Swedish Brain Foundation (FO2017-0075), Ming Wai Lau Centre for Reparative Medicine and Karolinska Institutet.

## Author contributions

KC/FP designed experiments with input from AMF/GCB. KC performed and analyzed all experiments, with input from AMF/GCB. KC/KZ/RH performed and analyzed qPCR experiments and IHC. KC/IK performed and analyzed EAE experiments. KC/EE/MJ analyzed RNA-seq. KC/FP wrote the manuscript with input from co-authors.

---

**Fig. S1.**
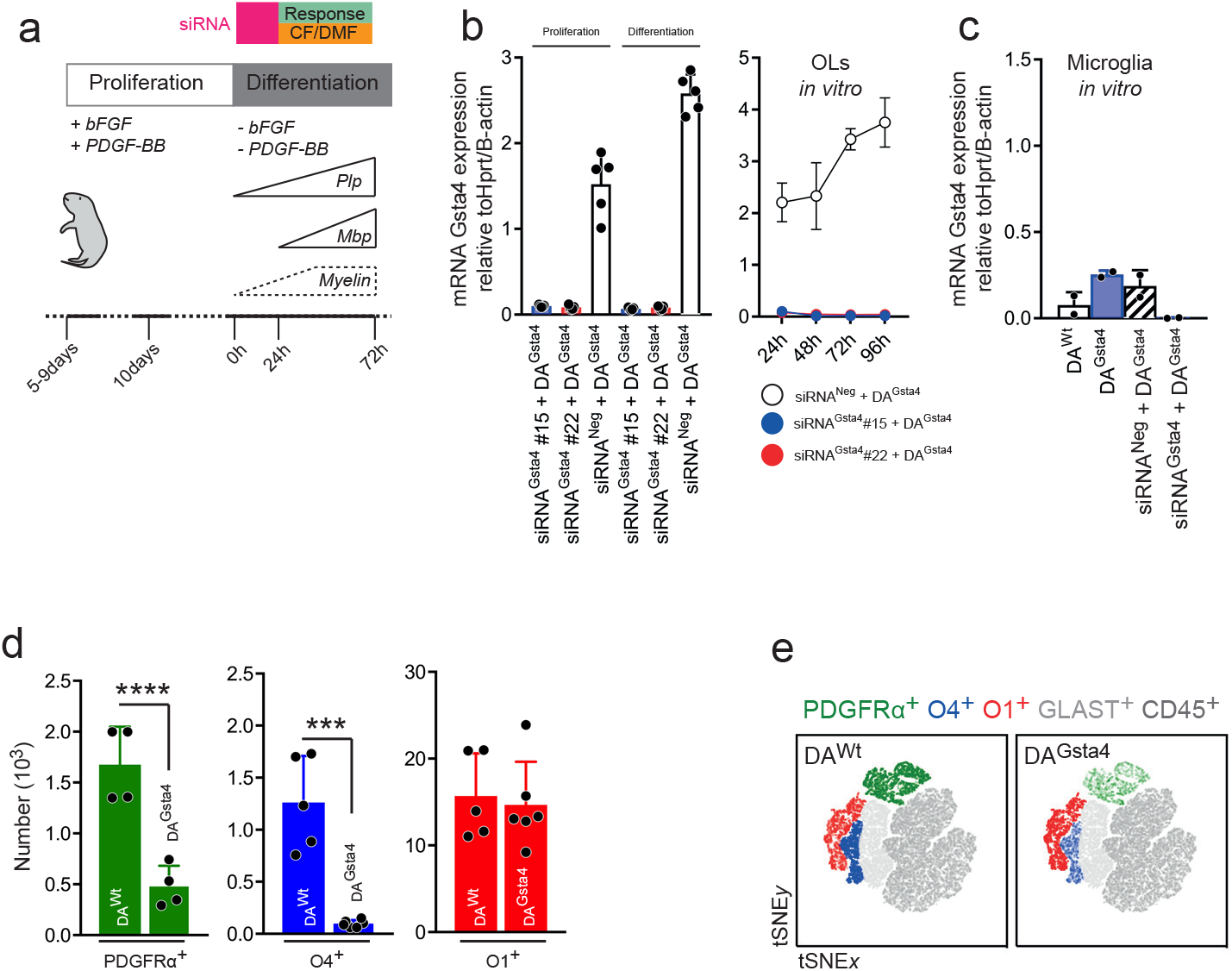
(a) Illustration of experimental set up for cell cultures. (b *left*) Transcription of *Gsta4* in proliferating OPCs and differentiated OLs 24h following 24h of incubation with siRNA^Gsta4^#15 siRNA^Gsta4^#22, targeting two separate regions of Gsta4 mRNA, or siRNA^Neg^. (b *right*) OPCs incubated with indicated siRNAs for 24h following switch to differential media and analysis of Gsta4 mRNA, 24, 48, 72 and 96h after siRNA wash-out. (c) siRNA knockdown of Gsta4 in microglia analyzed 24h after siRNA wash-out. (d) Total number of PDGFRα^+^ OPCs *(green)* (n=4), O4^+^ *(blue)* (n=5+6) and O1^+^ *(red)* (n=5+6) OLs from adult CC analyzed with flow cytometry. (e) tSNE plot based on indicated markers from two representative animals shows the distribution of cells across the OL-lineage. Graphs show representative results from three (d), two (b, c). All graphs show mean and S.D‥ Analysis in (d) was performed with one-way ANOVA. \*\*\**P*<0.001, \*\*\*\**P*<0.0001.

**Fig. S2.**
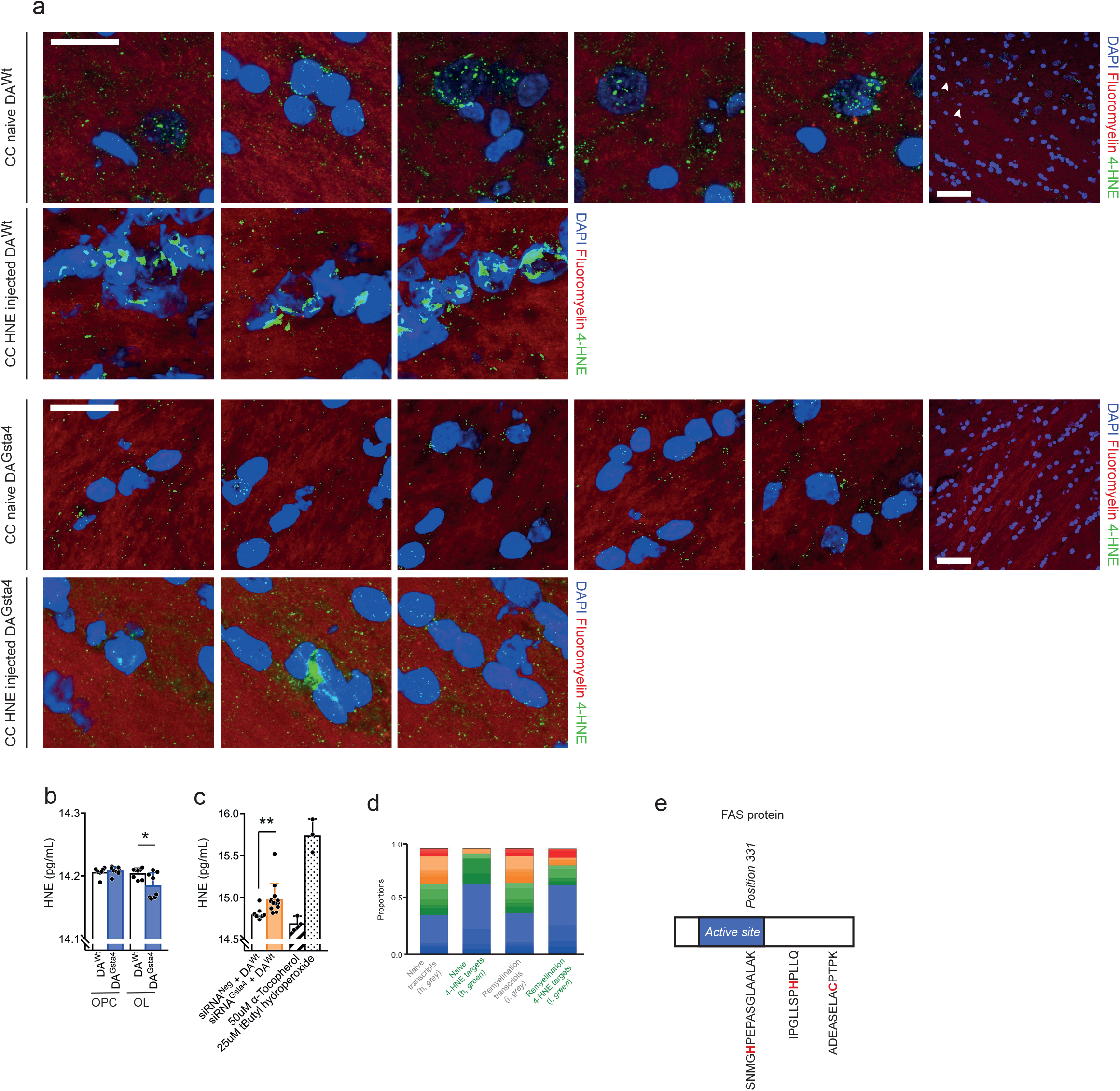
(a) Representative confocal images of anti-4-HNE staining of naïve CC from DA^Wt^ and DA^Gsta4^ and following intraparenchymal injections of HNE. (b) Intracellular 4-HNE load in cultured DA^Wt^ and DA^Gsta4^ OLs and OPCs cultures (n=8). (c) Intracellular 4-HNE load in cultured OLs following Gsta4 knockdown with siRNA-Gsta4 (n=14). (d) Porportions of subcellular location of proteins potentially modified with 4-HNE. (e) schematic illustration of Fas protein with three possible binding site for 4-HNE^60^. Experiment (b, c) shows two pooled experiments. All data are shown as mean and S.D. Student’s two-tailed unpaired *t* test was used for (d, e). \**P*<0.05,\*\**P* <0.01.

**Fig. S3.**
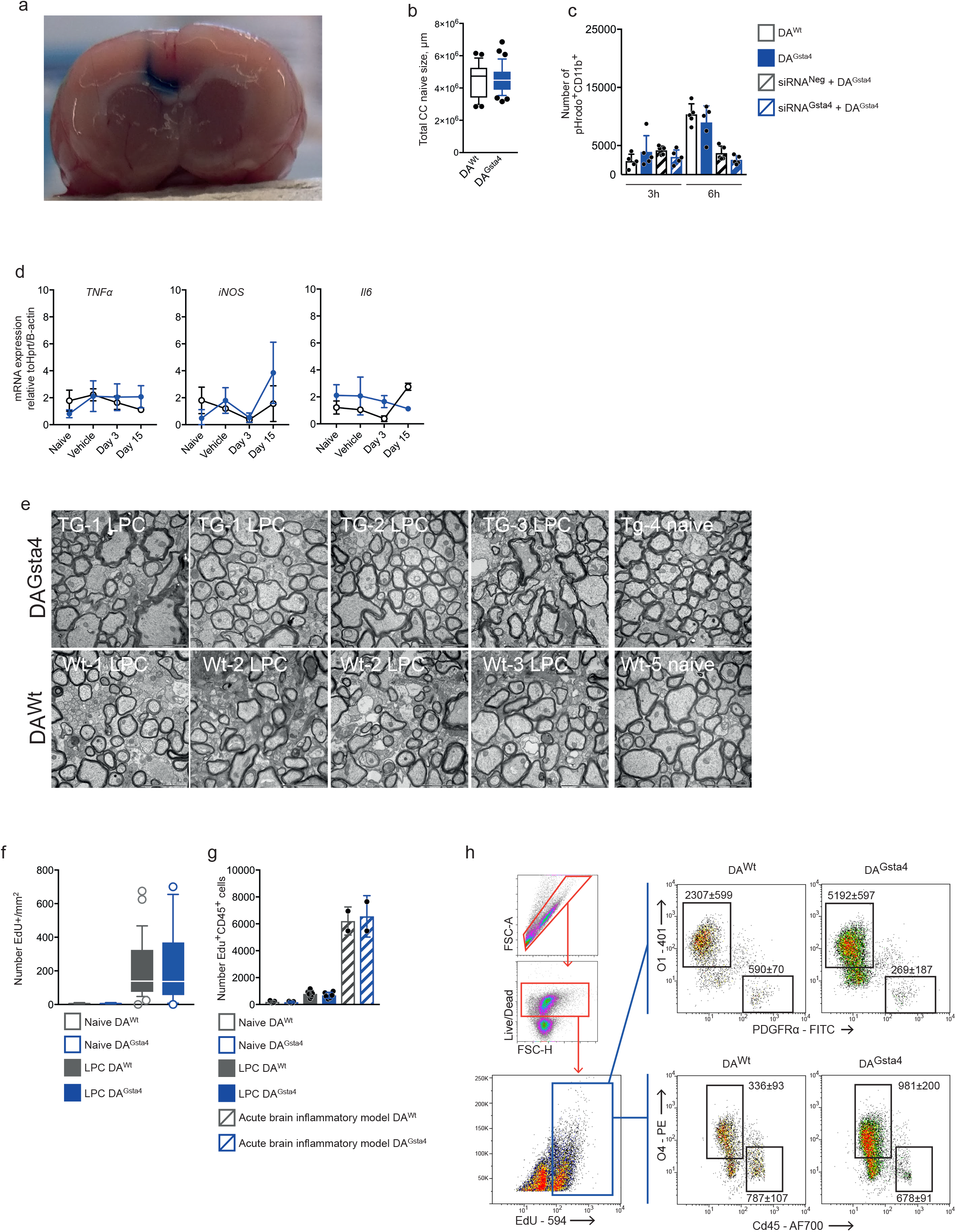
(a) Injection of blue dye due to the stereotactic coordinates used for LPC/HNE injection and stated in the method. (b) Total area of naïve corpus callosum between DA^Wt^ and DA^Gsta4^. (c) Evaluation of phagocytosis in primary microglia from DA^Wt^ or DA^Gsta4^, without (n=5) or with (n=5) knockdown of Gsta4 with siRNA-Gsta4. (d). Transcription of TNFα, iNos and Il6 in bulk CC tissue (n=3+3) (e). Representative TEM images from naïve DA^Wt^ and DA^Gsta4^ and following LPC (f). Total number of EdU labeled cells in CC evaluated by immunofluorescence. (g) Number of double-positive CD45^+^EdU^+^ cells ten days following LPC injections or application of traumatic brain injury model. Experiment (c) was repeated twice. All graph show mean and S.D., apart from (b, f) showing box and whiskers also indicating values outside 5-95 percentile.

**Fig. S4.**
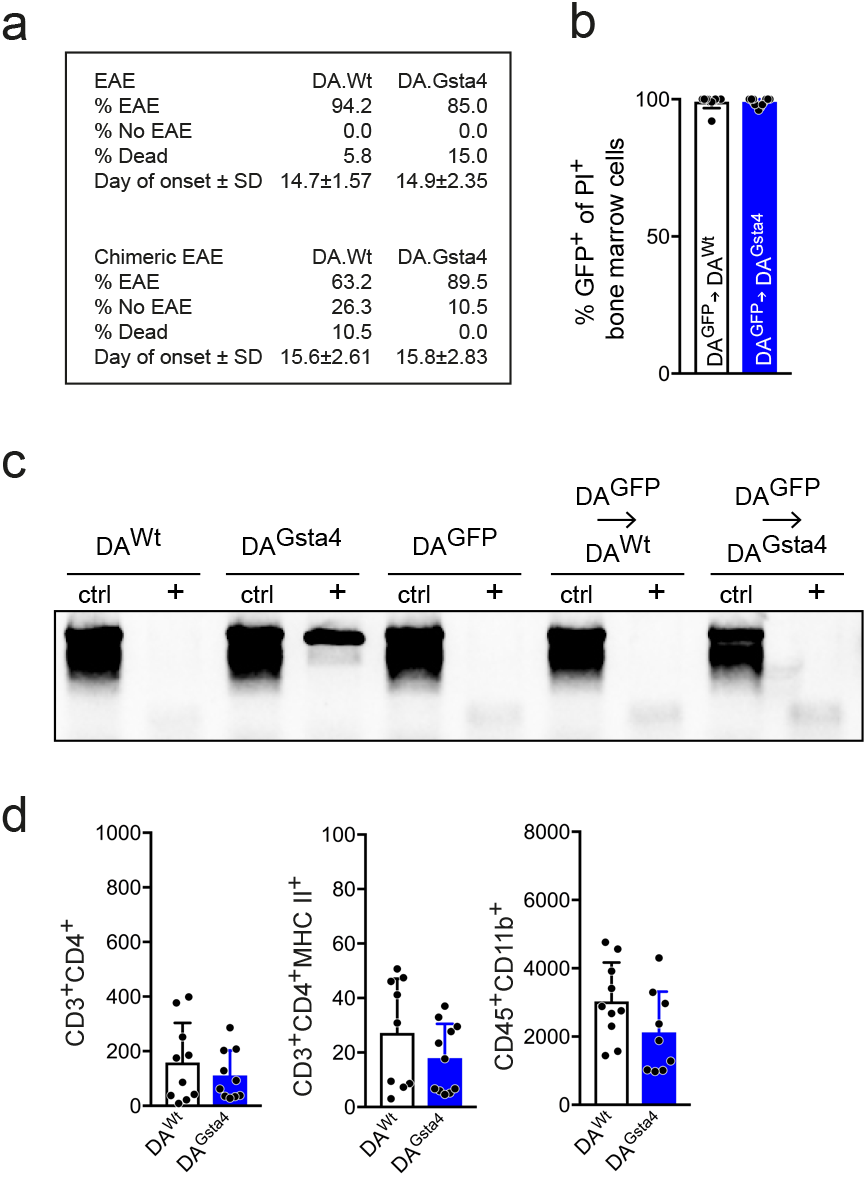
(a) EAE statistics. (b) Percent GFP^+^ of all propidium iodide^+^ bone marrow cells at the time of MOG immunization (n=12+16). (c) Representative genotype data of peripheral blood at the day of EAE termination. (d) Levels of T cells (CD3^+^CD4^+^) activated T cells (CD3^+^CD4^+^MHC II^+^) and microglia (CD45^+^CD11b^+^) in spinal cord from EAE experiment (n=9). All graphs shows mean and S.D.

## Notes

### Competing Interest Statement

The authors have declared no competing interest.

## References

1. Fünfschilling, U. et al. Glycolytic oligodendrocytes maintain myelin and long-term axonal integrity. Nature 485, 517–521 (2012).

2. Lee, Y. et al. Oligodendroglia metabolically support axons and contribute to neurodegeneration. Nature 487, 443–448 (2012).

3. Larson, V. A. et al. Oligodendrocytes control potassium accumulation in white matter and seizure susceptibility. Elife 7, 11298 (2018).

4. Nave, K.-A. & Werner, H. B. Myelination of the nervous system: mechanisms and functions. Annu. Rev. Cell Dev. Biol. 30, 503–533 (2014).

5. Prineas, J. W., Kwon, E. E., Cho, E. S. & Sharer, L. R. Continual breakdown and regeneration of myelin in progressive multiple sclerosis plaques. Ann. N. Y. Acad. Sci. 436, 11–32 (1984).

6. Ffrench-Constant, C. & Raff, M. C. Proliferating bipotential glial progenitor cells in adult rat optic nerve. Nature 319, 499–502 (1986).

7. Moyon, S. et al. Demyelination causes adult CNS progenitors to revert to an immature state and express immune cues that support their migration. J. Neurosci. 35, 4–20 (2015).

8. Smith, K. J., Blakemore, W. F. & McDonald, W. I. Central remyelination restores secure conduction. Nature 280, 395–396 (1979).

9. Ghosh, A. et al. Targeted ablation of oligodendrocytes triggers axonal damage. PLoS ONE 6, e22735 (2011).

10. Wolswijk, G. Chronic stage multiple sclerosis lesions contain a relatively quiescent population of oligodendrocyte precursor cells. J. Neurosci. 18, 601–609 (1998).

11. Jeffery, N. D. & Blakemore, W. F. Locomotor deficits induced by experimental spinal cord demyelination are abolished by spontaneous remyelination. Brain 120 (Pt 1), 27–37 (1997).

12. Chang, A., Nishiyama, A., Peterson, J., Prineas, J. & Trapp, B. D. NG2-positive oligodendrocyte progenitor cells in adult human brain and multiple sclerosis lesions. J. Neurosci. 20, 6404–6412 (2000).

13. Dutta, R. & Trapp, B. D. Mechanisms of neuronal dysfunction and degeneration in multiple sclerosis. Prog. Neurobiol. 93, 1–12 (2011).

14. Pohl, H. B. F. et al. Genetically induced adult oligodendrocyte cell death is associated with poor myelin clearance, reduced remyelination, and axonal damage. J. Neurosci. 31, 1069–1080 (2011).

15. Teunissen, C. E., Dijkstra, C. & Polman, C. Biological markers in CSF and blood for axonal degeneration in multiple sclerosis. Lancet Neurol 4, 32–41 (2005).

16. Barres, B. A. & Raff, M. C. Control of oligodendrocyte number in the developing rat optic nerve. Neuron 12, 935–942 (1994).

17. Trapp, B. D., Nishiyama, A., Cheng, D. & Macklin, W. Differentiation and death of premyelinating oligodendrocytes in developing rodent brain. J. Cell Biol. 137, 459–468 (1997).

18. Butts, B. D., Houde, C. & Mehmet, H. Maturation-dependent sensitivity of oligodendrocyte lineage cells to apoptosis: implications for normal development and disease. Cell Death Differ. 15, 1178–1186 (2008).

19. Sun, L. O. et al. Spatiotemporal Control of CNS Myelination by Oligodendrocyte Programmed Cell Death through the TFEB-PUMA Axis. Cell 175, 1811–1826.e21 (2018).

20. Prineas, J. W., Barnard, R. O., Kwon, E. E., Sharer, L. R. & Cho, E. S. Multiple sclerosis: remyelination of nascent lesions. Ann. Neurol. 33, 137–151 (1993).

21. Tripathi, R. B., Rivers, L. E., Young, K. M., Jamen, F. & Richardson, W. D. NG2 glia generate new oligodendrocytes but few astrocytes in a murine experimental autoimmune encephalomyelitis model of demyelinating disease. J. Neurosci. 30, 16383–16390 (2010).

22. Duncan, I. D. et al. The adult oligodendrocyte can participate in remyelination. Proc. Natl. Acad. Sci. U.S.A. 115, E11807–E11816 (2018).

23. Hill, R. A., Li, A. M. & Grutzendler, J. Lifelong cortical myelin plasticity and age-related degeneration in the live mammalian brain. Nat. Neurosci. 21, 683–695 (2018).

24. Barres, B. A., Jacobson, M. D., Schmid, R., Sendtner, M. & Raff, M. C. Does oligodendrocyte survival depend on axons? Curr. Biol. 3, 489–497 (1993).

25. Nave, K.-A. Myelination and the trophic support of long axons. Nat. Rev. Neurosci. 11, 275–283 (2010).

26. Almeida, R. & Lyons, D. Oligodendrocyte Development in the Absence of Their Target Axons In Vivo. PLoS ONE 11, e0164432 (2016).

27. Penderis, J., Shields, S. A. & Franklin, R. J. M. Impaired remyelination and depletion of oligodendrocyte progenitors does not occur following repeated episodes of focal demyelination in the rat central nervous system. Brain 126, 1382–1391 (2003).

28. Sim, F. J., Zhao, C., Penderis, J. & Franklin, R. J. M. The age-related decrease in CNS remyelination efficiency is attributable to an impairment of both oligodendrocyte progenitor recruitment and differentiation. J. Neurosci. 22, 2451–2459 (2002).

29. Doucette, J. R., Jiao, R. & Nazarali, A. J. Age-related and cuprizone-induced changes in myelin and transcription factor gene expression and in oligodendrocyte cell densities in the rostral corpus callosum of mice. Cell. Mol. Neurobiol. 30, 607–629 (2010).

30. Back, S. A. et al. Selective vulnerability of late oligodendrocyte progenitors to hypoxia-ischemia. J. Neurosci. 22, 455–463 (2002).

31. Baud, O. et al. Glutathione peroxidase-catalase cooperativity is required for resistance to hydrogen peroxide by mature rat oligodendrocytes. J. Neurosci. 24, 1531–1540 (2004).

32. Fragoso, G. et al. Developmental differences in HO-induced oligodendrocyte cell death: role of glutathione, mitogen-activated protein kinases and caspase 3. J. Neurochem. 90, 392–404 (2004).

33. Anderson, E. J., Katunga, L. A. & Willis, M. S. Mitochondria as a source and target of lipid peroxidation products in healthy and diseased heart. Clin. Exp. Pharmacol. Physiol. 39, 179–193 (2012).

34. Mahad, D., Ziabreva, I., Lassmann, H. & Turnbull, D. Mitochondrial defects in acute multiple sclerosis lesions. Brain 131, 1722–1735 (2008).

35. Lassmann, H., van Horssen, J. & Mahad, D. Progressive multiple sclerosis: pathology and pathogenesis. Nat Rev Neurol 8, 647–656 (2012).

36. Veiga, S., Ly, J., Chan, P. H., Bresnahan, J. C. & Beattie, M. S. SOD1 overexpression improves features of the oligodendrocyte precursor response in vitro. Neurosci. Lett. 503, 10–14 (2011).

37. Zhou, L., Zhang, H., Davies, K. J. A. & Forman, H. J. Aging-related decline in the induction of Nrf2-regulated antioxidant genes in human bronchial epithelial cells. Redox Biol 14, 35–40 (2018).

38. Kubben, N. et al. Repression of the Antioxidant NRF2 Pathway in Premature Aging. Cell 165, 1361–1374 (2016).

39. Haynes, R. L., Folkerth, R. D., Szweda, L. I., Volpe, J. J. & Kinney, H. C. Lipid peroxidation during human cerebral myelination. J. Neuropathol. Exp. Neurol. 65, 894–904 (2006).

40. Hemdan, S. & Almazan, G. Iron contributes to dopamine-induced toxicity in oligodendrocyte progenitors. Neuropathol. Appl. Neurobiol. 32, 428–440 (2006).

41. Thorburne, S. K. & Juurlink, B. H. Low glutathione and high iron govern the susceptibility of oligodendroglial precursors to oxidative stress. J. Neurochem. 67, 1014–1022 (1996).

42. Ćurko-Cofek, B. et al. Chronic iron overload induces gender-dependent changes in iron homeostasis, lipid peroxidation and clinical course of experimental autoimmune encephalomyelitis. Neurotoxicology 57, 1–12 (2016).

43. Esterbauer, H., Schaur, R. J. & Zollner, H. Chemistry and biochemistry of 4-hydroxynonenal, malonaldehyde and related aldehydes. Free Radic. Biol. Med. 11, 81–128 (1991).

44. Yang, Y., Huycke, M. M., Herman, T. S. & Wang, X. Glutathione S-transferase alpha 4 induction by activator protein 1 in colorectal cancer. Oncogene 35, 5795–5806 (2016).

45. Leonarduzzi, G., Robbesyn, F. & Poli, G. Signaling kinases modulated by 4-hydroxynonenal. Free Radic. Biol. Med. 37, 1694–1702 (2004).

46. Schneider, C., Porter, N. A. & Brash, A. R. Routes to 4-hydroxynonenal: fundamental issues in the mechanisms of lipid peroxidation. J. Biol. Chem. 283, 15539–15543 (2008).

47. Picklo, M. J., Amarnath, V., McIntyre, J. O., Graham, D. G. & Montine, T. J. 4-Hydroxy-2(E)-nonenal inhibits CNS mitochondrial respiration at multiple sites. J. Neurochem. 72, 1617–1624 (1999).

48. Linker, R. A. et al. Fumaric acid esters exert neuroprotective effects in neuroinflammation via activation of the Nrf2 antioxidant pathway. Brain 134, 678–692 (2011).

49. Scannevin, R. H. et al. Fumarates promote cytoprotection of central nervous system cells against oxidative stress via the nuclear factor (erythroid-derived 2)-like 2 pathway. J. Pharmacol. Exp. Ther. 341, 274–284 (2012).

50. Carlström, K. E. et al. Therapeutic efficacy of dimethyl fumarate in relapsing-remitting multiple sclerosis associates with ROS pathway in monocytes. Nat Commun 10, 3081 (2019).

51. Carlström, K. E. et al. Characterization of More Selective Central Nervous System Nrf2-Activating Novel Vinyl Sulfoximine Compounds Compared to Dimethyl Fumarate. Neurotherapeutics 14, 76–11 (2020).

52. Hirotsu, Y. et al. Nrf2-MafG heterodimers contribute globally to antioxidant and metabolic networks. Nucleic Acids Res. 40, 10228–10239 (2012).

53. Malhotra, D. et al. Global mapping of binding sites for Nrf2 identifies novel targets in cell survival response through ChIP-Seq profiling and network analysis. Nucleic Acids Res. 38, 5718–5734 (2010).

54. Mei, F. et al. Micropillar arrays as a high-throughput screening platform for therapeutics in multiple sclerosis. Nat. Med. 20, 954–960 (2014).

55. Cohen, J. A. & Tesar, P. J. Clemastine fumarate for promotion of optic nerve remyelination. Lancet 390, 2421–2422 (2017).

56. Green, A. J. et al. Clemastine fumarate as a remyelinating therapy for multiple sclerosis (ReBUILD): a randomised, controlled, double-blind, crossover trial. Lancet 390, 2481–2489 (2017).

57. Chandra, D. et al. Association of active caspase 8 with the mitochondrial membrane during apoptosis: potential roles in cleaving BAP31 and caspase 3 and mediating mitochondrion-endoplasmic reticulum cross talk in etoposide-induced cell death. Mol. Cell. Biol. 24, 6592–6607 (2004).

58. Zimniak, P. et al. Estimation of genomic complexity, heterologous expression, and enzymatic characterization of mouse glutathione S-transferase mGSTA4-4 (GST 5.7). J. Biol. Chem. 269, 992–1000 (1994).

59. Chen, Y. et al. Chemoproteomic profiling of targets of lipid-derived electrophiles by bioorthogonal aminooxy probe. Redox Biol 12, 712–718 (2017).

60. Yuan, W. et al. Highly Selective and Large Scale Mass Spectrometric Analysis of 4-Hydroxynonenal Modification via Fluorous Derivatization and Fluorous Solid-Phase Extraction. Anal. Chem. 89, 3093–3100 (2017).

61. Goebbels, S. et al. Elevated phosphatidylinositol 3,4,5-trisphosphate in glia triggers cell-autonomous membrane wrapping and myelination. J. Neurosci. 30, 8953–8964 (2010).

62. Fancy, S. P. J. et al. Dysregulation of the Wnt pathway inhibits timely myelination and remyelination in the mammalian CNS. Genes Dev. 23, 1571–1585 (2009).

63. Huang, J. K. et al. Retinoid X receptor gamma signaling accelerates CNS remyelination. Nat. Neurosci. 14, 45–53 (2011).

64. Marques, S. et al. Oligodendrocyte heterogeneity in the mouse juvenile and adult central nervous system. Science 352, 1326–1329 (2016).

65. Ineichen, B. V., Zhu, K. & Carlström, K. E. Axonal mitochondria across species adjust in diameter depending on thickness of surrounding myelin. bioRxiv 850370 (2018).

66. Ontaneda, D., Thompson, A. J., Fox, R. J. & Cohen, J. A. Progressive multiple sclerosis: prospects for disease therapy, repair, and restoration of function. Lancet 389, 1357–1366 (2017).

67. Gregson, N. A. Lysolipids and membrane damage: lysolecithin and its interaction with myelin. Biochem. Soc. Trans. 17, 280–283 (1989).

68. Ousman, S. S. & David, S. Lysophosphatidylcholine induces rapid recruitment and activation of macrophages in the adult mouse spinal cord. Glia 30, 92–104 (2000).

